# Promiscuous and multivalent interactions between Eps15 and partner protein Dab2 generate a complex interaction network

**DOI:** 10.1101/2024.09.30.615448

**Authors:** Andromachi Papagiannoula, Ida Marie Vedel, Kathrin Motzny, Maud Tengo, Arbesa Saiti, Sigrid Milles

## Abstract

Clathrin-mediated endocytosis depends on complex protein interactions. Eps15 plays a key role through interactions of its three EH domains with Asn-Pro-Phe (NPF) motifs in intrinsically disordered regions (IDRs) of other endocytic proteins. Using nuclear magnetic resonance spectroscopy, we investigate the interaction between Eps15’s EH domains and a highly disordered Dab2 fragment (Dab2₃₂₀₋₄₉₅). We find that the EH domains exhibit binding promiscuity, recognizing not only the NPF motif of Dab2 but also other phenylalanine containing motifs. This promiscuity enables interactions with Eps15’s own IDR (Eps15_IDR_), which lacks NPF motifs, suggesting an self-inhibitory state that promotes liquid-liquid phase separation. Despite competing for the same EH domain binding sites, Eps15_IDR_ and Dab2₃₂₀₋₄₉₅ can bind EH123 simultaneously, forming a highly dynamic interaction network that facilitates the recruitment of Dab2₃₂₀₋₄₉₅ into Eps15 condensates. Our findings provide molecular insights into the competitive interactions shaping the early stages of clathrin-mediated endocytosis.

## Introduction

Clathrin-mediated endocytosis (CME) is the major pathway for cargo uptake into the eukaryotic cell often with transmembrane receptors as cargoes. The important cellular process requires a complex network of interactions, which finally results in the formation and uptake of a clathrin coated vesicle. During the early phases of the process that takes roughly one to two minutes^1^, intrinsically disordered protein regions (IDRs) of the endocytic initiators play a major role ^2^. While folded proteins mainly exist in one stable conformation, IDRs inter-convert rapidly between many different conformations due to their relatively flat energy landscape. IDRs often recognize their folded partner proteins through small interaction regions of only a few residues, also called linear motifs^3,4^. This is also the case in CME, where IDRs of endocytic initiators interact with, for example, the major adaptor protein complex AP2 or clathrin^5–9^, allowing them to establish a robust multivalent interaction network which allows for rapid rearrangement of the different protein members. Eps15, together with FCHo1/2 proteins, counts among the first proteins to arrive at the endocytic pit^10^. It has three N-terminal Eps15 homology domains (EH domains), which share the common fold of two helix-loop-helix motifs (EF-hands) spaced by a short β-sheet^11^. The EH domains are followed by a central coiled-coil domain, through which Eps15 dimerizes^12,13^, and a large IDR of around 400 residues in length (Fig. 1A)^14^. Eps15 tetramers, formed by two anti-parallel Eps15 dimers have also been observed, seemingly requiring interaction between the EH domains and the IDR from opposite dimers^13^. FCHo1/2, which also forms dimers, use their C-terminal µ homology domain (µHD) to interact with the multiple Asp-Pro-Phe (DPF) motifs contained in Eps15’s IDR^15^. The multivalent interaction network between Eps15 and FCHo1/2 enables formation of liquid-like protein droplets, which has been proposed as a process to catalyze the early steps of CME^16^. The EH domains of Eps15, for example, can bind to Asn-Pro-Phe (NPF) motif containing proteins^17^, such as Epsin1, Stonin2^18^ or Dab2 (disabled homolog 2)^19^. While interactions between EH domains and both Epsin1 and Stonin2 have been characterized at molecular detail, the nature of Dab2’s interaction with EH domains remains to date elusive. Dab2 has an N-terminal folded phosphotyrosine interacting domain (PID), which it uses to anchor itself to the membrane and transmembrane cargo, followed by a large IDR of around 500 residues in length, which comprises both DPF as well as NPF motifs (Fig. 1B)^20^. Both Eps15 and Dab2 belong to the class of clathrin associated sorting proteins (CLASPs), which regulate cargo sorting during the early phases of CME and usually rely on downstream factors, such as the major adaptor protein complex AP2 or clathrin^3,21,22^. Dab2 has special properties as a CLASP since it can lead to successful CME of specific cargoes even in the absence of AP2. These cargoes, for example integrin β1, are specifically recognized by the Dab2 PID. Importantly, binding of Eps15 to Dab2 is required for the AP2 independent internalization of these protein cargoes^19^, highlighting the importance of understanding the communication between Eps15 and Dab2.

**Figure 1.**
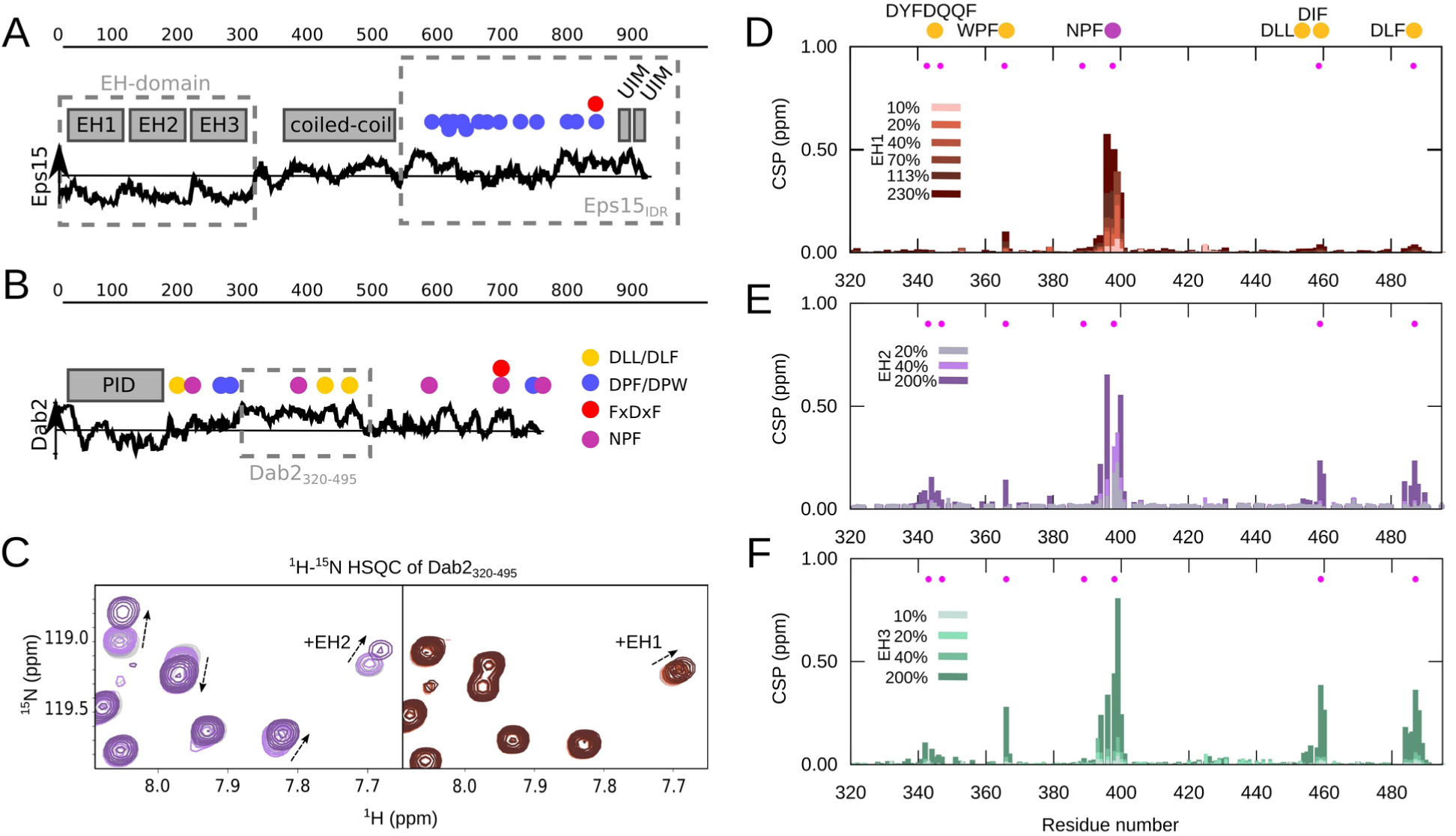
Interaction between Dab2_320-495_ and the individual EH domains of Eps15. Schematic illustration of **(A)** Eps15 and **(B)** Dab2. The disorder prediction (IUPred2A^23^) is shown as a black curve along the sequence. Above horizontal line: predicted disordered, Below horizontal line: predicted ordered. The folded domains are shown as gray boxes above the disorder prediction. Putative interaction motifs are shown as colored circles along the sequence (legend on the right of Dab2). DLL/DLF: clathrin binding; DPF/DPW: AP2α binding; FxDxF: AP2β binding; NPF: EH domain binding. The stretches used in this study are boxed with gray dashed lines. **(C)** Zoom into a ^1^H-^15^N HSQC spectrum of Dab2_320-495_ in the absence and presence of EH2 (left) and EH1 (right). **(D) – (F)** CSPs of 100 μM Dab2_320-495_ in the presence versus the absence of increasing concentrations of EH1 (D), EH2 (E) and EH3 (F), respectively. Color legends are displayed in the respective plots. Filled pink circles denote positions of phenylalanines.

Using nuclear magnetic resonance (NMR) spectroscopy, we investigate the region of Dab2 that is predicted to be most disordered, residues 320-495 (Dab2_320-495_), its conformational sampling, as well as its interactions with the EH domains of Eps15. All EH domains interact preferentially with the only NPF motif within Dab2_320-495_, but EH2 and EH3 demonstrate a high level of binding promiscuity towards other phenylalanine containing motifs. This promiscuity is maintained for the combined EH domains, comprising EH1, EH2 and EH3 (EH123), and surprisingly enables Eps15’s own IDR (Eps15_IDR_), which does not possess any NPF motifs, to bind to EH123 and thereby partially occupy the binding pockets of the different EH domains. We provide a detailed molecular characterization of these intra-molecular interactions within Eps15, which are likely key to liquid-liquid phase separation of Eps15 in the absence of other endocytic proteins. Our NMR results indicate that both Dab2_320-495_ and Eps15_IDR_ can bind EH123 simultaneously, but partially compete with each other. In line with this, we observe that Dab2_320-495_ is recruited into liquid-like condensates formed by full length Eps15. Our results provide a detailed description of the multivalent and promiscuous interactions between Eps15 and Dab2, which appear to be key to a dynamic protein network that can phase separate and evolve over time, which is required in the early stages of CME.

## Results

### Dab2_320-495_ is an IDR with mild helical propensities

The around 500 residue long IDR of Dab2 comprises a central region, which is predicted by IUPred2A^23^ to be significantly more disordered than the remainder of the IDR (Fig. 1B). We designed a construct encompassing this region, reaching from residue 320 to 495 (Dab2_320-495_). This region contains 2 putative binding sites for clathrin (DLL/DLF) and one NPF motif, prone to bind to EH domains contained in Eps15. We expressed and purified the construct, and assigned the NMR backbone resonances of a ^1^H, ^15^N, ^13^C labeled sample (Supplementary Fig. 1). We were initially not able to assign two small regions within Dab2_320-495_, for which we designed two new stretches (Dab2_328-360_ and Dab2_358-390_), whose ^1^H-^15^N heteronuclear single quantum coherence (HSQC) spectrum overlaid well with the one of Dab2_320-495_ (Supplementary Fig. 2), enabling the transfer of assignments from Dab2_328-360_ and Dab2_358-390_ to Dab2_320-495._ As predicted, Dab2_320-495_ appears to be disordered along its entire length, testified by secondary chemical shifts (SCSs) close to zero (Supplementary Fig. 3). However, calculating a conformational ensemble on the basis of chemical shifts using a combination of the statistical coil generator flexible meccano^24^ and the selection algorithm ASTEROIDS^25^ revealed three regions with mild helical propensity as compared to random coil: 339-354, 454-460, and 486-490 (Supplementary Fig. 3). Around the NPF motif, no particular structural propensity could be observed. In agreement with this, ^15^N spin relaxation rates (R_1ρ_, R_1_, {^1^H}-^15^N heteronuclear Overhauser effect - hetNOE) are reminiscent of those of an intrinsically disordered protein (Supplementary Fig. 4), with increased rigidity, testified by increased R_1ρ_ and hetNOE rates, apparent around the transient helix between residues 328 and 360 (helix_N_). This helix is connected with the remainder of the disordered chain by a few amino acids of more rapid mobility. The more C-terminal transient helices are also characterized by slightly increased R_1ρ_ and hetNOE rates, although less pronounced than those of helix_N_.

### EH1 binds selectively to NPF motifs, while EH2 and EH3 are promiscuous binders

We then titrated the different Eps15 EH domains (EH1, EH2, or EH3) into ^15^N Dab2_320-495_ and recorded ^1^H-^15^N HSQC spectra at the different titration steps. While small chemical shift perturbations (CSPs) are observed very locally upon addition of EH1, the spectrum of Dab2_320-495_ is more significantly perturbed when EH2 and EH3 are present at comparable concentrations (Figs. 1C-F, Supplementary Figs. 5, 6). Plotting the CSPs of Dab2_320-495_ upon interaction with the different EH domains along its sequence (Figs. 1D-F) clearly illustrates that EH1, EH2, and EH3 have different binding patterns: EH1 is the most selective of all Eps15 EH domains, binding mainly to the NPF motif of ^15^N Dab2_320-495,_ while very small CSPs occur around residue F366 only at 200% EH1. EH2 and EH3, on the other hand, were able to interact with multiple small stretches on Dab2_320-495_, revealing a high degree of binding promiscuity. This binding behavior is also mirrored by decreased peak intensities (Supplementary Fig. 6) and increased R_1ρ_ rates around the respective interaction sites (Figs. 2A, B). Since a spin-lock field of 1500 Hz was used in the R_1ρ_ experiments, effectively quenching contributions of intermediate exchange to the relaxation rates, the observed increases in R_1ρ_ reflect the slowed rotational tumbling times of the interacting residues when in contact with the respective EH domain. Residues in between the interaction sites have R_1ρ_ rates similar to those of the unbound Dab2_320-495_, suggesting that the different interaction sites act independently from each other. Even though the largest CSPs were observed around the NPF motif when EH2 and EH3 were added, also residues around DYF, WPF, DLF, and DIF motifs were in contact with those two EH domains at higher EH:Dab2 ratios. Common to all motifs is the phenylalanine residue, suggesting an importance for this residue type in the binding to EH2 and EH3 (Figs. 1E, F). Interestingly, most of these motifs are located in the regions with increased helical propensity of Dab2_320-495_ (Supplementary Fig. 3).

**Figure 2.**
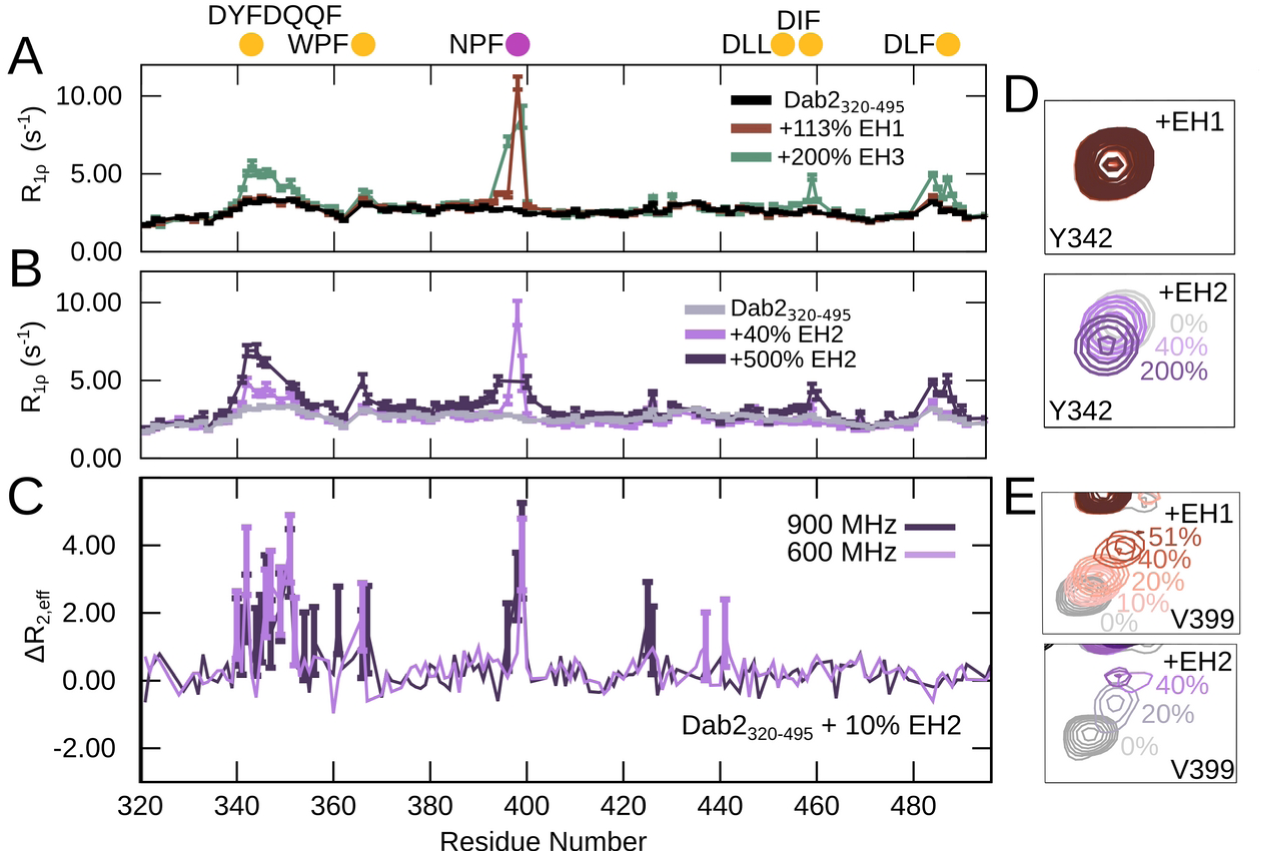
^15^N spin relaxation of Dab2_320-495_ upon interaction with the EH domains. ^15^N R_1ρ_ spin relaxation of Dab2_320-495_ (100 µM) in the absence and presence of **(A)** EH1 or EH3 and **(B)** increasing concentrations of EH2. The experiments were recorded at a ^1^H frequency of 600 MHz. **(C)** ΔR_2,eff_ calculated from effective R_2_ values (R_2,eff_) of relaxation dispersion experiments at a CPMG frequency of 31 Hz subtracted with R_2,eff_ at a CPMG frequency of 1000 Hz recorded on a sample of 100 µM Dab2_320-495_ in the presence of 10% EH2 at a ^1^H frequency of 600 MHz and 900 MHz. Error bars, propagated from the errors of R_2,eff_ at 31 Hz and 1000 Hz, are shown for values that are significantly larger than 0 (compared to the error). **(D)** Zoom into the peak corresponding to Y342 from Dab2_320-495_ ^1^H-^15^N HSQC spectra alone and in the presence of increasing concentrations of EH1 (top) and EH2 (bottom). **(E)** Zoom into the peak corresponding to V399 from Dab2_320-495_ ^1^H-^15^N HSQC spectra alone and in the presence of increasing concentrations of EH1 (top) and EH2 (bottom). Color legends are displayed in the respective panels.

The Dab2_320-495_ motifs were furthermore observed to bind to the EH domains with different dynamics. For example, V399, just next to the NPF motif, showed only mild peak broadening upon interaction with EH1, justified by the slowed rotational tumbling times when in complex. Upon interaction with EH2, however, significantly stronger broadening was observed (Fig. 2D, E). The other interacting regions show very little broadening and are mainly characterized by CSPs. These differential behaviors prompted us to record Carr-Purcell-Meiboom-Gill (CPMG) relaxation dispersion of Dab2_320-495_ to investigate the possibility of intermediate exchange in the microsecond to millisecond regime between the bound and unbound states in the presence of EH2 (Fig. 2C). These experiments confirm that the NPF motif binds EH2 in intermediate exchange on the NMR chemical shift timescale. Surprisingly, many residues throughout the helix_N_ also displayed dynamics on the intermediate timescale upon binding. The remaining motifs were concluded to bind in fast exchange.

In order to quantify the affinity of the interaction between Dab2_320-495_ and the EH domains, we attempted to extract dissociation constants (K_D_) for the different Dab2_320-495_ interaction motifs. Even though fitting of CSPs as a function of EH domain concentration was difficult due to the low affinities and peak broadening (especially around the NPF motif upon interaction of EH2), they can be estimated to be in the high micromolar to millimolar range (Supplementary Fig. 7A). Analysis of the CPMG relaxation dispersion curves of Dab2_320-495_ upon interaction with EH2 revealed a K_D_ value in a similar range (196 μM), whereby the dispersion in the helix_N_ and around the NPF motif were fit together (Supplementary Fig. 7B).

### Dab2_320-495_ binds to the main hydrophobic pocket of EH1, EH2, and EH3 and a second binding pocket of EH2

We wondered whether differences in the EH domains could explain the distinct interactions observed with the different motifs of Dab2_320-495_. We thus set out to determine the binding pockets on each of the EH domains that engage in interaction with Dab2_320-495_. Even though NMR assignments of the three EH domains exist^26–28^, we assigned all three EH domains under our experimental conditions (Supplementary Figs. 8-10). From the carbon SCSs, we calculated secondary structure propensities (SSPs)^29^ which are in agreement with previously determined NMR structures of the proteins (Supplementary Fig. 11)^30–32^. Additionally, ^15^N R_1_ and R_1ρ_ relaxation rates reflect small folded domains with flexible N- and C-termini as expected (Supplementary Fig. 12). We then titrated Dab2_320-495_ into the different ^15^N labeled EH domains and recorded ^1^H-^15^N Transverse relaxation-optimized spectroscopy (TROSY) spectra of the EH domains at the different titration steps (Figs 3). The largest CSPs can be observed around the previously identified binding pockets^18,31^ and are comparable between all three EH domains (Fig. 3D). An additional binding region, previously shown to bind a second NPF motif in Stonin2^18^, is clearly visible in EH2. The region seems to participate in binding also within EH1 and EH3, albeit in a less pronounced way (Fig. 3). R_1ρ_ relaxation rates systematically increase throughout all residues when Dab2_320-495_ is titrated into the ^15^N EH domains, likely reflecting mildly increased tumbling times resulting from the motional drag that the bound IDR exerts (Fig. 3F)^33^.

**Figure 3.**
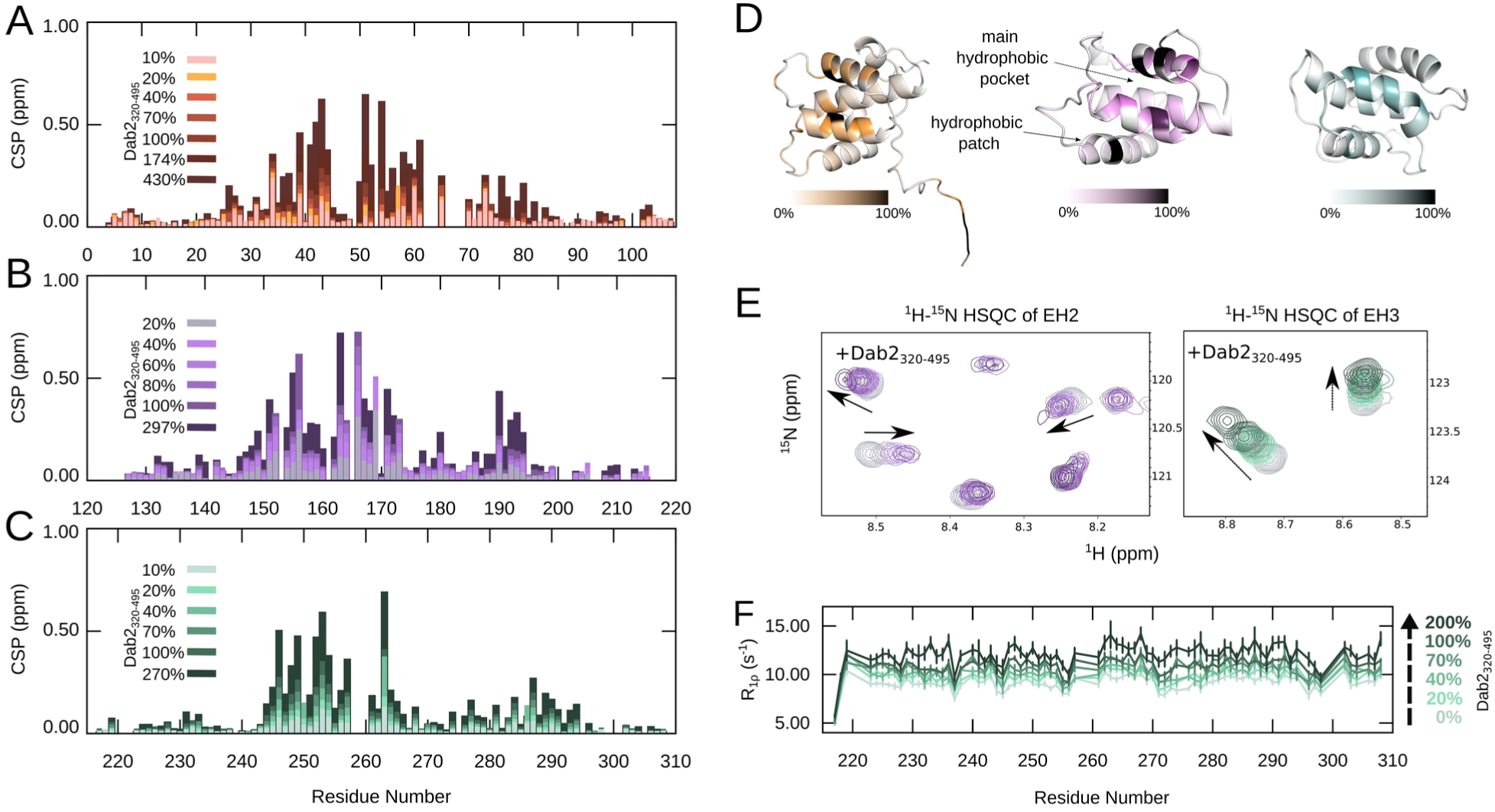
Interaction between the individual EH domains of Eps15 and Dab2_320-495_. **(A-C)** CSPs of 100 μM EH1 (A), EH2 (B) and EH3 (C), respectively, in the absence versus the presence of increasing amounts of Dab2_320-495_. Color legends are displayed in the respective plots. **(D)** Structures of EH1 (AlphaFold2^34^), EH2 (PDB 1FF1^31^) and EH3 (PDB 1C07^32^) with the CSP at 100% Dab2_320-495_ mapped onto them. The linear gradient starts from 0% (white), which represents no CSP, to the 100% (black), which is 0.75 ppm (the biggest CSP observed between EH2 and Dab2_320-495_). **(E)** Zoom into a ^1^H-^15^N HSQC spectrum of EH2 (left) and EH3 (right) alone and in the presence of Dab2_320-495_. **(F)** ^15^N R_1ρ_ spin relaxation of EH3 (100 µM) in the absence and presence of Dab2_320-495_ at a ^1^H frequency of 600 MHz. Color legends are indicated in the figures.

### The full EH-domain containing region (EH123) shows increased binding promiscuity

Since the different EH domains are normally not isolated, but expressed in row in the context of Eps15, we investigated the interaction between Dab2_320-495_ and the full EH-domain containing region (EH1-EH2-EH3, also called EH123) by titrating increasing concentrations of EH123 into ^15^N Dab2_320-495_. As expected from the interactions of the individual EH domains, both CSPs and R_1ρ_ rates reveal that EH123 is able to interact with the same regions of Dab2_320-495_ as observed for EH2 and EH3 (Figs. 4A, B). Interestingly, the R_1ρ_ relaxation rates of Dab2_320-495_ in the presence of EH123 show increased relaxation not only of the residues of the five Dab2_320-495_ binding regions, but the residues located in between the binding motifs also show mildly increased rates. This is particularly evident for the residues located between the first three binding motifs (DYFDQQF, WPF and NPF) and suggests an overall stiffening of the chain, potentially due to binding of the motifs to different EH domains within the same EH123 molecule.

**Figure 4.**
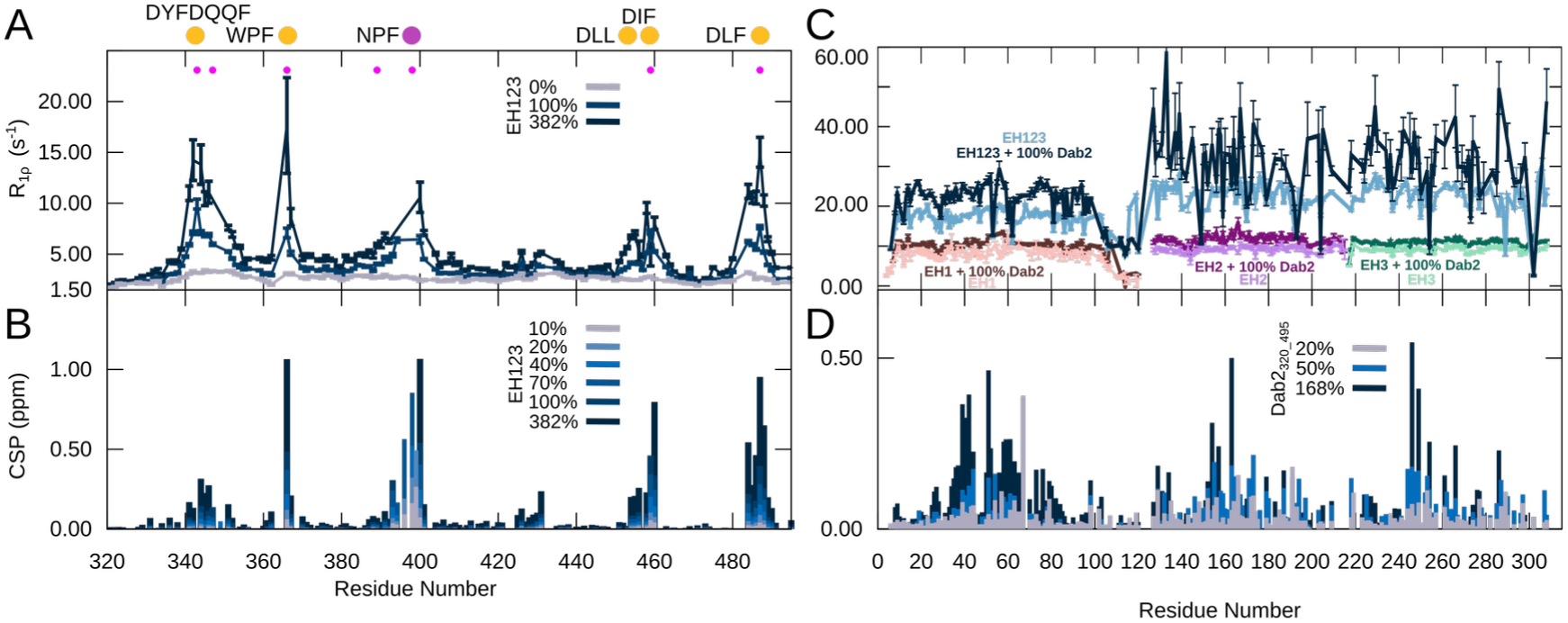
Interaction between EH123 and Dab2_320-495_. **(A)** ^15^N R_1ρ_ spin relaxation of Dab2_320-495_ (100 µM) in the absence and presence of increasing concentrations of EH123 at a ^1^H frequency of 600 MHz. **(B)** CSPs between the ^1^H-^15^N HSQC of ^15^N Dab2_320-495_ in the absence and presence of increasing amounts of EH123. The filled pink dots represent the phenylalanines present in the Dab2_320-495_. **(C)** ^15^N R_1ρ_ spin relaxation of the individual ^15^N EH domains (EH1, EH2, EH3) as well as EH123 (200 µM) in the absence and presence of 100% Dab2_320-495_ at a ^1^H frequency of 600 MHz. **(D)** CSPs between the ^1^H-^15^N TROSY of ^15^N EH123 in the absence and presence of increasing amounts of the Dab2_320-495_. The color legends are indicated in the respective figure panels.

While CSPs around the NPF motif remain the largest upon interaction with EH123, the difference between the interaction of EH123 with the NPF motif and the other motifs appears smaller than for interaction with the individual EH domains. Indeed, R_1ρ_ rates increase much more significantly around these promiscuous sites, albeit part of the increase should be attributed to the larger size of EH123 relative to the individual domains. Nonetheless, binding affinities approximated from the CSPs for EH123 binding are significantly higher than those of the individual EH domains (Supplementary Fig. 7A). This suggests that once one EH domain within EH123 is bound to Dab2_320-495_, the two other EH domains of the same molecule remain available for (preferential) binding to other motifs, an effect reminiscent of avidity, effectively increasing the total level of binding promiscuity. A much higher concentration of individual EH domains (3x as many) is needed to partially recover the chemical shift differences and R_1ρ_ rates of Dab2_320-495_ in the presence of EH123 (Supplementary Fig. 13).

### All EH domains within EH123 engage in interaction with Dab2

To investigate whether Dab2_320-495_ binds to the same sites on the EH domains when they are expressed in row rather than independently, we also aimed to characterize the interaction between Dab2_320-495_ and EH123 from the side of EH123. We therefore produced ^15^N labeled EH123 and recorded a ^1^H-^15^N TROSY spectrum. If the different EH domains were unaffected by being connected to each other, we would expect this spectrum to overlay perfectly with those of the individual EH domains. However, small CSPs were observed for all three EH domains, and the peaks of EH2 and EH3 appeared broadened (Supplementary Fig. 14), revealing that each EH domain is affected by the presence of the two additional domains potentially due to intra-molecular interactions. Despite the changes in the spectrum of EH123 compared to those of EH1, EH2, and EH3, the assignments from the individual domains could be fully transferred to the combined construct. In order to examine the broadening of the peaks originating from EH2 and EH3, we recorded ^15^N spin relaxation of EH123 (R_1ρ_ and hetNOEs). All R_1ρ_ rates of EH123 were significantly increased as compared to the individual EH domains which was expected for this larger protein construct (Fig. 4C, Supplementary Fig. 14C). The rates of EH2 and EH3, however, were significantly higher overall than those from EH1. Indeed, the increased R_1ρ_ rates were similar for EH2 and EH3, suggesting that EH2 and EH3 tumble together as one entity, while EH1 can move independently within EH123. HetNOEs, which are sensitive to motion in the picosecond range, and thus faster motion than R_1ρ_ rates, show quite similar values all along EH123, reflecting the similar fold and internal dynamics of the individual domains (Supplementary Fig. 14C). Only the linker between EH1 and EH2 displays much lower hetNOE values, demonstrating that this linker is undergoing much more rapid motion than the rest of the protein construct, also compared to the linker connecting EH2 and EH3, in very good agreement with EH2 and EH3 tumbling together within the EH123 construct.

When Dab2_320-495_ was added to ^15^N EH123, we observed CSPs for all three domains, similar to those of the individual EH domains when interacting with Dab2_320-495_ (Fig. 4D), suggesting that the binding sites on all EH domains are available also within EH123, despite the fact that EH2 and EH3 tumble together within EH123. R_1ρ_ rates increased throughout EH123 when Dab2_320-495_ was added, and the rates of EH1 remained lower than those of EH2 and EH3, suggesting that the overall conformation of the EH domains within EH123 is preserved upon binding of Dab2_320-495_.

### Eps15’s own IDR interacts with EH123

The remarkable binding promiscuity towards phenylalanine containing motifs observed between EH123 and Dab2_320-495_, prompted us to investigate whether EH123 could also interact with Eps15’s own IDR (Eps15_IDR_). Eps15_IDR_ does not contain any canonical NPF motifs, but contains 14 DPF motifs (Fig. 1A), which have previously been suggested to interact with Eps15’s own EH domains^16^. When we added equimolar amounts of Eps15_IDR_ to ^15^N EH123, CSPs were observed throughout the ^1^H-^15^N TROSY spectrum of EH123, confirming the interaction and further suggesting that all EH domains bind to Eps15_IDR_ (Fig. 5A, Supplementary Fig. 15). Upon interaction, most peaks of EH2 and EH3 were severely broadened, likely because of the motional drag the long Eps15_IDR_ exerts on EH123. As a consequence, many of the signals were too weak to calculate CSPs and extract relaxation rates for EH2 and EH3 within EH123 upon interaction with Eps15_IDR_. However, the CSPs originating from EH1 within EH123 were all smaller than those in presence of the same amount of Dab2_320-495_, indicating that EH1 interacts less well with Eps15_IDR_ than with Dab2_320-495_, in agreement with EH1 being the least promiscuous of the domains, binding more selectively to NPF motifs than EH2 and EH3 (Fig. 5A).

**Figure 5.**
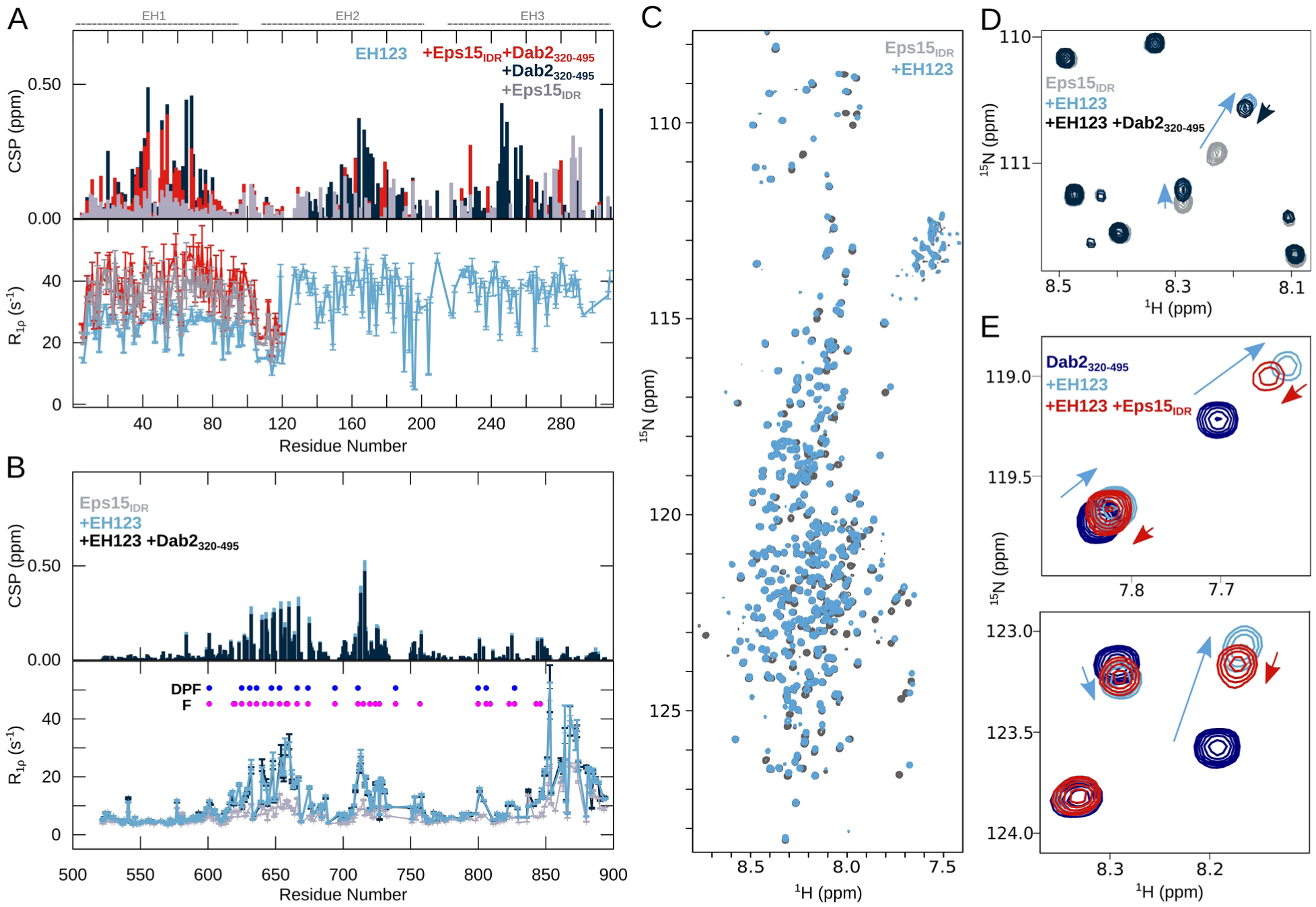
Competitive binding of Dab2_320-495_ and Eps15_IDR_ to EH123. **(A)** CSPs calculated between the ^1^H-^15^N TROSY of 200 µM ^15^N EH123 in the absence and presence of 200 µM Dab2_320-495_ and/or 200 µM Eps15_IDR_ and ^15^N R_1ρ_ spin relaxation of ^15^N EH123 in the absence and presence of 100% Dab2_320-495_ and/or 100% Eps15_IDR_ at a ^1^H frequency of 1200 MHz. Only the R_1ρ_ rates of EH1 in the presence of Dab2_320-495_/Eps15_IDR_ are shown as the peaks of EH2 and EH3 were severely broadened. The parts of EH123 corresponding to EH1, EH2, and EH3 are illustrated above the plots. **(B)** CSPs calculated between the ^1^H-^15^N HSQC of 100 µM ^15^N Eps15_IDR_ in the absence and presence of 100 µM EH123 and 100 µM EH123 + 100 µM Dab2_320-495_ and ^15^N R_1ρ_ spin relaxation of ^15^N Eps15_IDR_ in the absence and presence of 100% EH123 and 100% EH123 + 100% Dab2_320-495_ at a ^1^H frequency of 1200 MHz. **(C)** ^1^H-^15^N HSQC spectra of 100 µM ^15^N Eps15_IDR_ alone and in the presence of 100 µM EH123. **(D)** Zoom into ^1^H-^15^N HSQC spectrum of 100 µM ^15^N Eps15_IDR_ alone and in the presence of 100 µM EH123 and 100 µM EH123 + 100 µM Dab2_320-495_ **(E)** Zoom into ^1^H-^15^N HSQC spectrum of 100 µM ^15^N Dab2_320-495_ alone and in the presence of 100 µM EH123 and 100 µM EH123 + 100 µM Eps15_IDR_. Color codes are denoted in the respective panels.

### EH123 interacts with DPF clusters and other phenylalanine containing motifs in Eps15_IDR_

In order to further characterize the interaction between Eps15_IDR_ and EH123, we assigned the backbone resonances of ^1^H, ^15^N, ^13^C labeled Eps15_IDR_. To circumvent the spectral overlap of the long Eps15_IDR_, we designed four overlapping smaller stretches (Eps15_IDR 481-581_, Eps15_IDR 569-671_, Eps15_IDR 648-780_, and Eps15_IDR 761-896_). The ^1^H-^15^N HSQC spectra of these stretches overlayed nicely with the spectrum of the full Eps15_IDR_ (Supplementary Fig. 16), allowing the transfer of ^1^H and ^15^N resonance assignments between the spectra. We assigned 242 out of 393 resonances distributed across the IDR except for the N-terminal residues 481-521 to which no resonances could be assigned (Supplementary Fig. 17). This is likely because these residues (481-504) are part of the coiled-coil domain as predicted by AlphaFold2^34^, giving rise to weak peaks in the 3D assignment spectra. Low R_1_ and R_1ρ_ relaxation rates and SCSs around 0 for both Eps15_IDR_ and the smaller stretches agree well with a disordered protein (Supplementary Fig. 18). The C-terminal part however (∼residues 850-885) has increased R_1ρ_ rates and SSP values close to 1, revealing a stable alpha helix. This alpha helix, which is also predicted by AlphaFold2^34^, contains the two ubiquitin interaction motifs (UIMs) present in Eps15^35,36^. In general, the R_1ρ_ relaxation rates are slightly larger within Eps15_IDR_ compared to the smaller stretches, potentially resulting from small hydrophobic clusters in these regions leading to transient self-interactions within the chain – an effect that has been observed for transiently folded elements in other intrinsically disordered proteins (IDPs)^37^.

We next recorded a ^1^H-^15^N HSQC spectra of ^15^N Eps15_IDR_ with equimolar amounts of unlabeled EH123 in order to pinpoint the exact interaction regions on Eps15_IDR_. R_1ρ_ rates, CSPs, and intensity ratios compared to Eps15_IDR_ on its own revealed two relatively large interaction regions (Figs. 5B, C, Supplementary Fig. 19). These regions, 620-680 and 700-730, are characterized by a high density of phenylalanines. While the first region is enriched in DPF motifs, which have previously been proposed to bind to EH123, the second region is characterized by fewer DPF motifs and contains multiple other small linear motifs with phenylalanine, underlining the promiscuity of the EH domains. Moreover, small CSPs and increases in R_1ρ_ rates are observed around three smaller regions indicating additional interaction sites for EH123: residues ∼800-809 and ∼820-827 containing DPF(s), and residues ∼843-846 containing phenylalanines. While increases in R_1ρ_ relaxation rates and decreases in intensity ratios in the C-terminal helix are also observed (Fig. 5B, Supplementary Fig. 19), the small CSPs accompanying these changes suggest that this is through binding to the first helical residues (843-846), which then affect the tumbling time of the entire alpha helical element.

### Dab2_320-495_ and Eps15_IDR_ partially compete for binding to EH123

The interaction observed between EH123 and Eps15_IDR_ suggests that intramolecular interactions within Eps15 may occupy Dab2 binding sites on the EH domains in the native context. We therefore conducted a competition experiment, acquiring a spectrum of ^15^N EH123 with equimolar amounts of both Eps15_IDR_ and Dab2_320-495_. The resulting CSPs of EH123 showed that in the presence of both Dab2_320-495_ and Eps15_IDR_, the CSPs of EH1 were similar to those in presence of Dab2 only and accompanied by slightly increased R_1ρ_ rates, while the peaks of EH2 and EH3 were broadened severely similar to when only Eps15_IDR_ was added (Fig. 5A). While it is difficult to disentangle the individual contributions of Dab2_320-495_ and Eps15_IDR_ binding to the spectral changes, this suggest that both IDRs may bind EH123 at the same time. To further investigate this, we recorded a ^1^H-^15^N HSQC spectrum of ^15^N Eps15_IDR_ and added first EH123 and then Dab2_320-495_. While most peaks were unaffected by addition of Dab2_320-495_, some peaks shifted very slightly back towards the unbound state of Eps15_IDR_ (Figs. 5B, D, Supplementary Fig. 19). This suggests that binding between Eps15_IDR_ and EH123 is largely unaffected by the presence of Dab2_320-495_. We then conducted the same competition experiment, but this time using ^15^N Dab2_320-495_. Again, when adding equimolar amounts of Eps15_IDR_ to ^15^N Dab2_320-495_ bound to EH123, the spectrum of Dab2_320-495_ was only slightly affected, with peaks from bound Dab2_320-495_ moving towards the unbound form (Fig. 5E, Supplementary Fig. 20), likely due to competition with Eps15_IDR_. To rule out that any of the observed spectral changes could originate from a potential interaction between Dab2_320-495_ and Eps15_IDR_, we recorded a ^1^H-^15^N HSQC spectrum of ^15^N Dab2_320-495_ with Eps15_IDR_, confirming that the two IDRs do not interact (Supplementary Fig. 21). Taken together, the competition experiments conducted from the side of each binding partner (EH123, Eps15_IDR_, and Dab2_320-495_) suggest that both Eps15_IDR_ and Dab2_320-495_ can bind to EH123 at the same time, even though they are competing for the same EH domain interaction sites. This could be possible due to a fast on and off rate of the low affinity interaction motifs, thereby creating a complex and dynamic interaction network allowing both IDRs to bind EH123.

### Dab2_320-495_ enters into Eps15 protein condensates

Eps15 has recently been shown to form liquid-like droplets, and interactions between the EH domains and Eps15_IDR_ were proposed to contribute to droplet formation^16^. The seemingly simultaneous interactions of Eps15_IDR_ and Dab2_320-495_ with EH123 observed here, made us question whether Dab2_320-495_ could be recruited to Eps15 condensates. We therefore expressed and purified full length Eps15, from here on just called Eps15, and labeled its cysteines with the fluorophore Alexa 594. In order to fluorescently label Dab2_320-495_, which does not contain any cysteines, we created a single cysteine mutant (S328C Dab2_320-495_), and labeled it with Alexa 488. We first examined if we could achieve liquid-like droplet formation of 7 µM Eps15 under our experimental conditions. Indeed, upon addition of 3% PEG, we observed droplet formation (Fig. 6A). Unlike Eps15, Dab2_320-495_ was not able to form any droplets on its own under the same conditions (Fig. 6B). When Eps15 and Dab2_320-495_ were mixed in ratios of 1:1 and 1:20 in the presence of 3% PEG, Dab2_320-495_ readily localized within the Eps15 condensates (Fig. 6C, D). This indicates, in line with our NMR results, that the intermolecular Eps15 interactions responsible for droplet formation are maintained in the presence of Dab2_320-495_, while Eps15 also remains available for interaction with Dab2.

**Figure 6.**
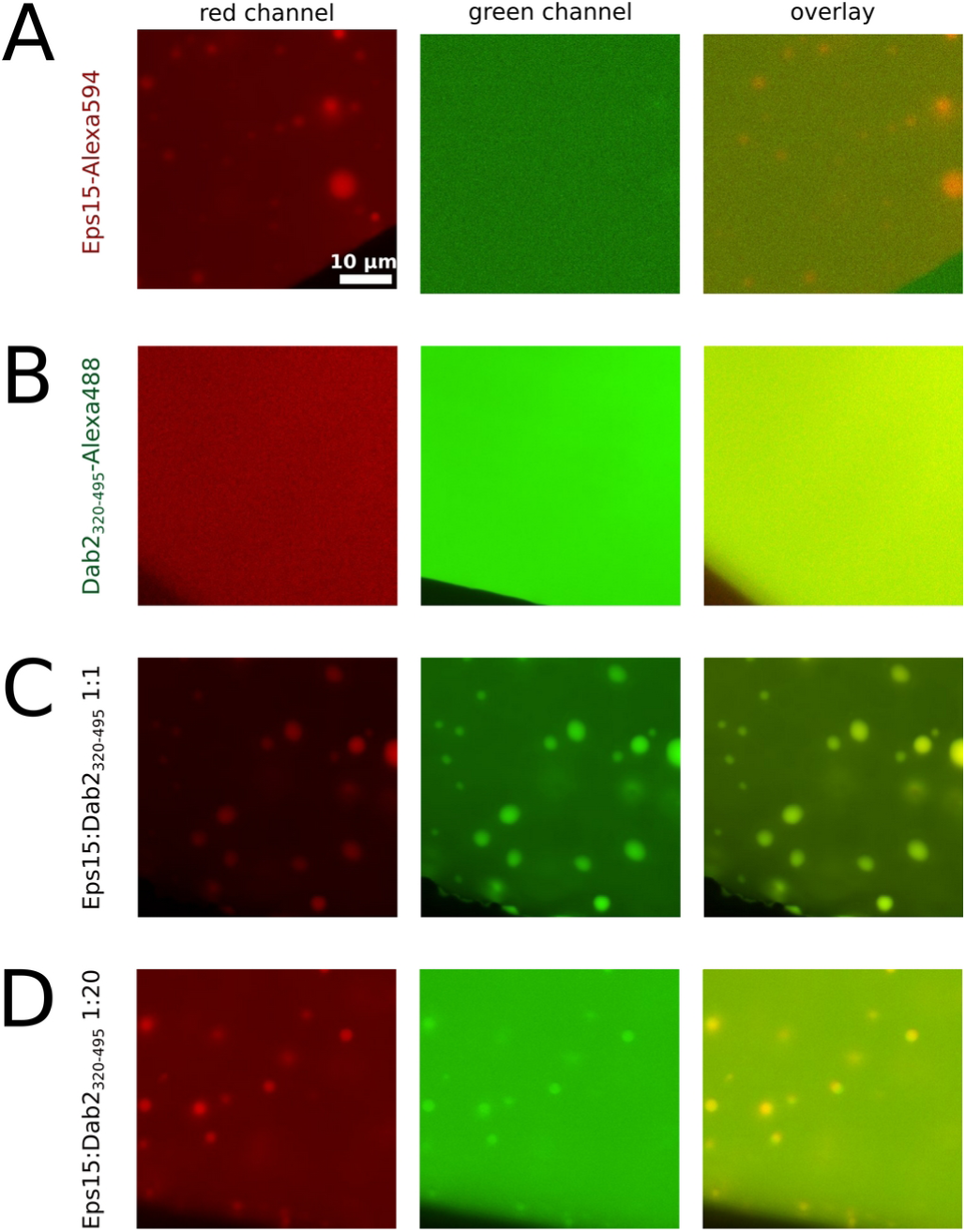
Dab2_320-495_ is recruited into Eps15 droplets. Microscopy images showing **(A)** droplets of Eps15 alone, **(B)** the absence of liquid-liquid phase separation of Dab2_320-495_ alone, and **(C,D)** Eps15 and Dab2_320-495_ in a 1:1 or 1:20 ratio showing recruitment of Dab2 into Eps15 droplets.

## Discussion

The dynamic network formed by intrinsically disordered regions (IDRs) and folded proteins interacting with short linear motifs within these IDRs, is key to a number of biological processes ^4,38^. Clathrin-mediated endocytosis is a prime example for a process that comprises many of those short linear motifs^2,21^, leading to the formation of a complex interaction network. Known examples are the α and β2 appendage domains of the major adaptor protein AP2 that interact with DPF/DPW and FxDxF motifs (x = any amino acid)^3^, the clathrin heavy chain terminal domain that interacts with DLL/DLF motifs^7,8^ or the EH domains found in Eps15, which interact with NPF motifs of diverse IDR partners^18^. Although extensive studies have identified these consensus sequences, it has meanwhile become clear that additional binding sites are yet to be discovered and that this requires a strategy by which focus lies on studying long endocytic IDRs. This has recently allowed identifying a particularly long interaction site between the neuronal AP180 and the AP2β2 appendage domain^7^. In addition, many endocytic IDRs seem to bind to the same partners^39^ and understanding how the different binding modes synergize or compete with each other is crucial for a molecular comprehension of clathrin-mediated endocytosis.

In this study, we shed light onto those partially competitive interactions by investigating binding of a large intrinsically disordered region stemming from the CLASP Dab2 (Dab2_320-495_) and its interactions with Eps15 EH domains, as well as its competition for binding with Eps15’s own IDR (Eps15_IDR_). In line with the previous literature and extensive peptide screens to assess binding patterns of Eps15 EH domains^40^, all three EH domains of Eps15 (EH1, EH2 and EH3) prefer to bind to the only NPF motif within the sequence of Dab2_320-495_ (Figs. 1, 2). In the presence of EH2, significant exchange in the microsecond to millisecond time scale is observed around the NPF motif and an N-terminal transient helix in Dab2_320-495_ (helix_N_), allowing to estimate a dissociation constant between Dab2_320-495_ (NPF motif and helix_N_) and EH2 on the order of hundreds of micromolar (Fig. 2, Supplementary Fig. 7). These affinities are in agreement with affinities of other small linear motif interactions determined in the context of clathrin-mediated endocytosis^7–9^, but orders of magnitudes weaker than those observed between Stonin2 and EH2 (K_D_ in the nanomolar range)^18^. Although the EH domains bind preferentially to the NPF motif of Dab2_320-495_, as it is expected from the literature^18,31,32,40^, the atomic resolution provided by our NMR experiments also points toward additional, slightly weaker binding sites including DYF, WPF, DLF, and DIF motifs (Fig. 1). Such sites will certainly play a role in the crowded environment of the endocytic pit, and these motifs further testify to an extremely promiscuous binding behavior of EH2 and EH3. This is in line with the observation that other EH domains, such as the EH domain of EHD1 or of POB-1 can bind to the motif xPF^41,42^. By investigating the EH domain interactions with small linear motifs from the side of a bona fide IDR, we can conclude that the promiscuity of EH2 and EH3 is even larger than previously anticipated, binding to most phenylalanine containing motifs (Figure 7A).

**Figure 7.**
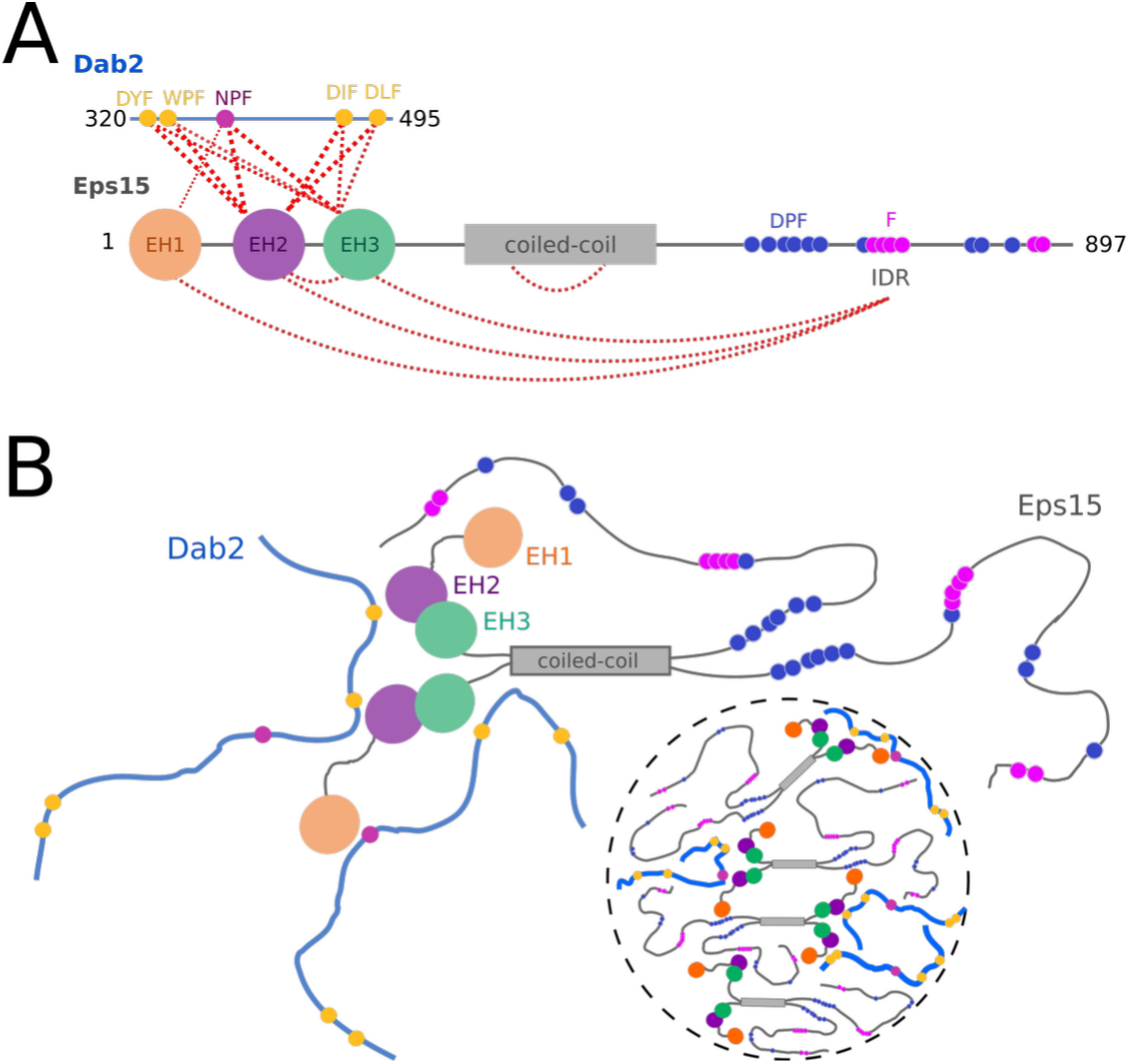
**(A)** Schematic illustration of the complex interaction network between Eps15 and Dab2_320-495_ including intramolecular interactions within Eps15. Interactions are visualized with red dashed lines. **(B)** Cartoon of self-interactions within Eps15 and of interactions between Eps15 and Dab2_320-495_. Eps15’s EH domains interact with Eps15_IDR_ and with Dab2_320-495_. EH2 and EH3 tumble together. All three EH domains interact with the NPF motif within Dab2_320-495_, while EH2 and EH3 are promiscuous binders. The weak multivalent interaction network of the two IDR’s with EH123 likely contribute to co-phase separation of Eps15 and Dab2_320-495_.

Even though some proteins contain individual EH domains, this is not the case for Eps15, which contains EH1, EH2 and EH3 sequentially at its N-terminus. Unexpectedly, we observe that EH2 and EH3 move together as one entity in a construct comprising all three EH domains (EH123, Fig. 4C), suggesting a potential interaction between EH2 and EH3. EH1, on the other hand, tumbles independently from the other two EH domains. Nonetheless, EH123 makes all EH domain binding sites available for interaction with Dab2_320-495_. From the side of Dab2_320-495_, binding to EH123 is strikingly different than binding to the individual EH domains. While the same NPF and promiscuous motif interactions maintain, the non-NPF interactions seem to gain importance, possibly enhanced through avidity effects of the three EH domains in close spatial proximity compared to the individual domains (Fig. 4, Supplementary Fig. 13).

The present promiscuity of interactions, particularly in the context of the full EH123 construct, suggests that the EH domains might bind to Eps15’s own IDR, which has also been suggested previously^13,16,42^. Addition of Eps15_IDR_ to ^15^N EH123 indeed led to CSPs and increased ^15^N R_1ρ_ relaxation rates confirming this intramolecular interaction (Fig. 5A). While this interaction has previously been proposed to occur through DPF motifs contained in Eps15_IDR_^16^, our data recorded on ^15^N labeled Eps15_IDR_ shows that EH123 binds to all phenylalanine containing regions of Eps15_IDR_ highlighting the promiscuity of EH123 also observed in the binding to Dab2_320-495_ (Fig. 5B). While one would expect that an NPF containing IDR, such as Dab2_320-495_, might replace Eps15_IDR_ from EH123 due to the preferred interaction of EH domains with NPF motifs compared to more promiscuous binding sites, this seems not to be the case. When both Eps15_IDR_ and Dab2_320-495_ are added to ^15^N labeled EH123, the spectrum and corresponding relaxation rates show signatures of binding to both proteins, although the individual contributions cannot be disentangled due to their similar interaction sites on EH123 (Fig. 5A). A similar behavior is observed when a sample containing all three proteins is detected from the side of ^15^N Eps15_IDR_ or ^15^N Dab2_320-495_. Only few regions within the spectra of Eps15_IDR_ and Dab2_320-495_ slightly revert towards their unbound state when both EH123 and the respective other IDR (Dab2_320-495_ or Eps15_IDR_) are added (Fig. 5D, E), suggesting that both Eps15_IDR_ and Dab2_320-495_ can bind EH123 at the same time.

How is this possible when Dab2_320-495_ and Eps15_IDR_ occupy the same binding sites on EH123? Key to this question is likely the multivalency by which both IDRs interact with EH123, such that, while some motifs of Dab2_320-495_ might be displaced from EH123 in a dynamic fashion when Eps15_IDR_ is added – or vice versa - others maintain, thereby creating a dynamic trimeric (or even higher order) complex. Which of the motifs remain bound to EH123 and which get displaced is certainly a function of the individual motif’s affinities. This is easiest visualized as a dynamic equilibrium where motifs in Dab2 are competing with Eps15’s own IDR for binding to the EH domains, forming a complex and dynamic network (Fig. 7). The fact that neither Dab2_320-495_ or Eps15_IDR_ can effectively outcompete the other, suggests that the strengths of the individual interactions between EH123 and Dab2_320-495_ or Eps15_IDR_ are not vastly different. This further argues for a physiological relevance of the interactions seen between the EH domains and Eps15_IDR_ and the competition effects observed.

While binding of EH domains to an intrinsically disordered linker of the same protein has been observed for the EH domain contained in EHD2^43^, the interaction network established by Eps15 is certainly remarkable, since so many interactions take place and compete with each other (Fig. 7). This has implications for all other binding partners of Eps15, whether binding to the EH domains or Eps15’s IDR, since any binding partner will be competing with these intramolecular interactions. Indeed, the interactions observed between Eps15 EH domains and Eps15_IDR_ could constitute some kind of auto-inhibitory mechanism, such as recently described for WW domain proteins^44^, which could be (partially) released by other interaction partners, such as Dab2. In this context it should be noted, that the interactions between EH123 and Eps15_IDR_ could be both intramolecular within one Eps15 molecule and intermolecular between different Eps15 molecules. Indeed, Eps15-Eps15 interactions have previously been suggested to drive Eps15 phase separation in the context of clathrin-mediated endocytosis. For example, based on previous literature on the interaction between the EH domain of POB1 and DPF motifs^42^, which are also contained in Eps15_IDR_, interaction between Eps15 EH domains and Eps15_IDR_ have been suggested to be important for liquid-liquid phase separation^16^. While our molecular data show that not only DPF motifs, but also other F-rich protein regions are involved in this interaction, other interactions of similar strengths between Eps15 and its binding partners, may drive this partner into the liquid-like droplets, such as it has been observed for FCHo1/2^16^ or ubiquitin^45^. Of note, phenylalanines have been identified as critical ‘stickers’ to promote weak interactions in liquid-liquid phase separation of IDPs^46^. Usually, they are thought to interact with phenylalanines in other IDPs, thereby creating a dynamic interaction network. In the case of Eps15, we have observed phenylalanines to contact small folded domains (EH domains) – interactions that may also drive Dab2_320-495_ into Eps15 droplets. In good agreement with this hypothesis, we observe that Dab2_320-495_, which does not form droplets on its own, also enters liquid-like droplets of Eps15, likely due to the weak multivalent network between the two (Figure 7B). Therefore, while current literature point to Eps15 as the main initiator of condensate formation during the early phases of CME^16^ our data suggest that Dab2 may be recruited into such condensates also in the cellular context, illustrating the role of a complex, promiscuous and multivalent interaction network to recruit downstream adaptors in endocytosis. Indeed, it will be interesting to see, whether the interactions between EH domains and Eps15_IDR_ are maintained also in the presence of proteins containing multiple NPF motifs, binding with higher affinities^18^, and what consequences this has on liquid-liquid phase separation at the endocytic pit and thus progression of productive endocytosis.

## Materials and Methods

### Cloning

Eps15-pmCherryN1 was a gift from Christien Merrifield (Addgene plasmid # 27696 ; http://n2t.net/addgene:27696 ; RRID:Addgene_27696)^47^. The three individual EH domains (EH1, EH2, EH3) and Eps15_IDR 761-896_ were cloned into the pET41c vector leading to constructs with a non-cleavable C-terminal His-tag. Full length Eps15, EH123, Eps15_IDR_, Eps15_IDR 481-581_, Eps15_IDR 569-671_ and Eps15_IDR 648-780_ were cloned into the pET28a vector with an N-terminal GB1 solubility tag pET28-6His-GB1. This leads to the expression of a TEV (tobacco etch virus) cleavable 6His-GB1-TEV site construct followed by the protein of interest. The gene of Dab2_320-495_ was purchased from Twist Bioscience and cloned into a pET28a vector with non-cleavable C-terminal His-tag. For the purpose of fluorescence labeling, a single cysteine mutant of Dab2_320-495_ was constructed using site-directed mutagenesis (Dab2_320-495 S328C_). Dab2_328-360_ and Dab2_358-390_ were cloned into a pET28a vector with an N-terminal GB1 solubility tag pET28-6His-GB1. The UniProt IDs of Eps15 and Dab2 used in this study are P42566 and P98082, respectively.

### Protein expression and purification

The proteins were expressed in the *E. coli* Rosetta (DE3) strain and grown in LB medium with 30 mg/L Kanamycin and 30 mg/L Chloramphenicol at 37°C. When the optical density (OD) at 600 nm was around 0.6-1, expression was induced with 1 mM isopropyl-β-D-thiogalactopyranoside (IPTG) and the expression was then continued at 20°C overnight. For isotope labeling (^15^N, ^13^C) M9 minimal medium was used and supplemented with 1g/L ^15^NH_4_Cl and/or with 2 g/L ^13^C-glucose. Cells were lysed by sonication in lysis buffer (20 mM Tris, 150 mM NaCl pH 8, with Roche Ethylenediaminetetraacetic Acid (EDTA)-free protease inhibitor cocktail (Sigma-Aldrich Chemie GmbH)). Purification involved a two-step process: initial nickel purification followed by size-exclusion chromatography (SEC). The nickel column was equilibrated in lysis buffer before the filtered lysate was applied to a nickel column. The column was then washed with lysis buffer containing 20 mM imidazole, and the protein was eluted using lysis buffer supplemented with 400 mM imidazole. For all constructs expressed in the pET28-6His-GB1 vector, the eluted fraction, which contained the target protein (validated by SDS-PAGE and Coomassie staining), underwent overnight dialysis with 1 mg TEV protease at 4°C in 500 mL of lysis buffer supplemented with 5 mM β-mercaptoethanol before proceeding to the SEC purification step. Proteins were further purified using a Superdex 75 or a Superdex 200 column, equilibrated in NMR buffer (50 mM Na-phosphate pH 6, 150 mM NaCl, and 2 mM dithiothreitol (DTT)). Fractions containing pure protein (validated by SDS-PAGE) were concentrated and frozen with final protein concentrations determined by absorbance at 280nm using extinction coefficients determined by Expasy Protparam^48^.

### NMR spectroscopy

NMR experiments were measured at the Leibniz-Forschungsinstitut für Molekulare Pharmakologie (FMP), Berlin, Germany (^1^H frequencies of 600, 750, 900, 1200 MHz), and at the Institute of Structural Biology (IBS), Grenoble, France (^1^H frequencies of 600 MHz). All experiments were measured in NMR buffer (50 mM Na-phosphate pH 6, 150 mM NaCl, 2 mM DTT) at 25°C. Addition of up to 5 mM CaCl_2_ did not affect our results. Spectra were processed with NMRPipe^49^, using qMDD^50^ for non-uniformly sampled assignment spectra, and analyzed with CCPN^51^.

### ^15^N, ^13^C Backbone assignment

Almost complete assignments could be obtained for all protein constructs. The assignment spectra of the ^15^N, ^13^C labeled EH1, EH2, EH3, Dab2_328-360_, and Dab2_358-390_ were acquired at a ^1^H frequency of 750 MHz. Assignment spectra of Eps15_IDR_, Eps15_IDR 480-581_, Eps15_IDR 569-671_, Eps15_IDR 648-780_, Eps15_IDR 761-896_, and Dab2_320-495_ were acquired at a ^1^H frequency of 600 MHz. Standard BEST-TROSY triple resonance experiments correlating CO, Cα, Cβ resonances (HNCO, HNCOCA, HNCA, iHNCA, HNCOCACB, iHNCACB)^52^ were acquired. The assignments were done in CCPN^51^ and then validated using MARS^53^. Secondary chemical shifts and secondary structure propensities^29^ were calculated using random coil values from refDB^54^. Overlapping the 3 TROSY spectra of the individual EH domains (EH1, EH2, EH3) we transferred the ^1^H and ^15^N resonances to the TROSY of the EH123 domain. The assignment of the different Eps15_IDR_ stretches was transferred to the spectrum of the full Eps15_IDR_ in a similar way.

### Titrations, ^15^N relaxation, and relaxation dispersion

Extraction of peak intensities (I) as well as ^1^H and ^15^N chemical shifts, were carried out from ^1^H-^15^N HSQC or BEST-TROSY spectra. Combined chemical shift perturbations (CSPs) were calculated using

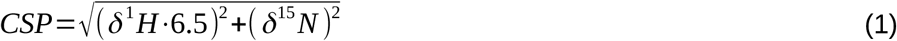

The specific concentrations used in the different titrations are indicated in the respective figures, with the percentage of partner indicating the molar ratio of partner protein:observed protein.

Residue specific *K*_d_ values were estimated by fitting the following equation to the CSPs as a function of concentration:

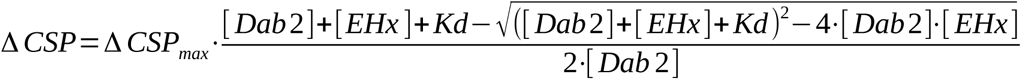

Fitting was performed with Python.

^15^N R_1ρ_, R_1_ and {^1^H}-^15^N HetNOE relaxation rates^55^ were assessed at 600 MHz and 1200 MHz ^1^H Larmor frequency. The spin-lock field for the R_1ρ_ experiment was set to 1500 Hz or 2000 Hz for the 600 MHz and 1200 MHz magnets, respectively, and 6-7 delays, between 10 and 230 ms (for the disordered Dab2_320-495_ and Eps15_IDR_) and between 10 and 70 ms (for the folded EH domains) were used to sample the decay of magnetization. R_1_ was measured using 6-7 delays between 0 to 1.2 s. Relaxation dispersion experiments^56^ were conducted at 600 and 900 MHz, employing 14 CPMG frequencies ranging from 31 to 1000 Hz, with a constant-time relaxation of 32 ms. Errors were determined based on standard deviations of repeat measurements. The minimum error was set to 0.5. For the experiment with 100 μM Dab2_320-495_ in the presence of 10% EH2, 12 residues within the helix_N_ and the NPF motif (Val340, Asp341, Gln345, Gln346, Gln349, Ser351, Thr354, Lys356, Phe366, Phe 398, Val399, Asp426) were fit globally using a 2 site exchange model with the software ChemEx (https://github.com/gbouvignies/chemex). An exchange rate of 149 ± 13 s^-1^ and a percentage bound of 3 ± 0.3% were obtained from the fit.

### Conformational ensemble of Dab2_320-495_

A conformational ensemble was calculated from Dab2_320-495_ and Dab2_328-360_ using a combination of the statistical coil generator flexible meccano^24^ and the genetic algorithm ASTEROIDS^25^. From a statistical coil ensemble of 10,000 conformers, 200 conformations that together best described the H_N_, N, CO, Cα and Cβ chemical shifts of the proteins were selected. A new ensemble of 8500 conformers was generated using the Φ and ψ angles from the previous selection and supplemented with 1500 conformations from the previous pool of conformers. A new selection of 200 conformations was performed on the new pool. This iteration was repeated 4 times. Ensemble averaged chemical shifts were generated using SPARTA^57^ and secondary chemical shifts were calculated based on RefDB^54^.

### Structures of the individual EH domains

The structures that were used to plot the CSPs for EH2 and EH3 are the PDB codes 1FF1^31^ and 1C07^32^, respectively. For EH1 we use an AlphaFold2^34,58^ prediction since at the time of writing no *human* Eps15 EH1 structure was deposited in the PDB. The CSPs of each EH domain with 100 μM Dab2_320-495_ (100%) were plotted onto the structures. The linear gradient is from 0%, which represents no CSP, up to 100%, which is the highest CSP observed at 0.75 ppm. The structures were visualized using Pymol^59^.

### Protein labeling with fluorescent dyes

Protein labeling was performed using malemeide dyes (Alexa 488 for Dab2_320-495 S328C_ and Alexa 594 for Eps15) essentially as described previously^59,60^. After adding 10 mM DTT overnight to fully reduce the proteins, they were dialyzed into a phosphate-buffer (50 mM Na-phosphate pH 7, 150 mM NaCl) for 2×1 hour at room temperature. At least 5 times molar excess of the dyes were used for the labeling reaction, which after mixing contained a maximum of 10% V/V DMSO. The protein/dye mixtures were allowed to react for 1 hour at room temperature, and incubated overnight at 4°C. To stop the reaction, 10 mM of DTT were added before injecting the mixtures on a SEC70 or 650 equilibrated in NMR buffer to remove any unconjugated dye.

### Protein droplets and microscopy imaging

All droplet formation assays were performed in a buffer of 50 mM Na-phosphate pH 6, 150 mM NaCl, and 2 mM DTT, 3% w/v PEG8000, using a total of 7 µM Eps15 with different ratios of Dab2_320-495_. For both proteins, a ratio of 100:1 of non labeled:fluorescently labeled protein was used. A total sample volume of 20 µL was placed in 8 wells polystrene chambers with 1.5 borosilicate coverglass (Nunc Lab-Tek). Epi-fluorescence imaging was performed with a Nikon Ti Eclipse microscope with a 60x (PLAN APO, NA:1.40, WD 0.13 mm) oil immersion objective. The setup was controlled by the imaging software NIS (Nikon). Three sequential images were taken per experiment. Microscopy images were imported and analyzed using ImageJ/Fiji^62^.

## Data and materials availability

All study data are included in the article and/or supporting information. The chemical shift assignments have been uploaded to the Biological Magnetic Resonance Data Bank (BMRB) under the accession numbers 52613 (Dab2_320-495_), 52866 (Eps15_IDR 481-581_), 52864 (Eps15_IDR 569-671_), 52863 (Eps15_IDR 648-780_), and 52867 (Eps15_IDR 761-896_), respectively.

## Acknowledgments

We thank all members of the Milles group for fruitful discussions and critical proofreading. We thank M.R. Jensen for providing the pET28-6His-GB1 plasmid, M. Blackledge and M.R. Jensen for providing NMR analysis scripts, P. Schmieder, N. Trieloff and M. Beerbaum for technical assistance on the NMR spectrometers and M. Lehmann from the Imaging Facility of FMP for technical support. The Institut de Biologie Structurale acknowledges integration into the Interdisciplinary Research Institute of Grenoble. This work was supported by the Leibniz-Forschungsinstitut für Molekulare Pharmakologie (FMP) (to S.M.). This project has received funding from the European Research Council (ERC) Starting Grant MultiMotif to S.M. under the European Union’s Horizon 2020 research and innovation program (grant agreement no. 802209).

## Author Contributions

SM, AP and IMV designed research. AP, IMV, KM, MT, AS performed research. AP, IMV, SM analyzed the data. AP, IMV, SM wrote the paper.

## Additional information

Supplementary information is available.

