## Supporting Information for "Promiscuous and multivalent interactions between Eps15 and partner protein Dab2 generate a complex interaction network"

Papagiannoula[#], Vedel[1#], *et al.*

[#]equal contribution

*

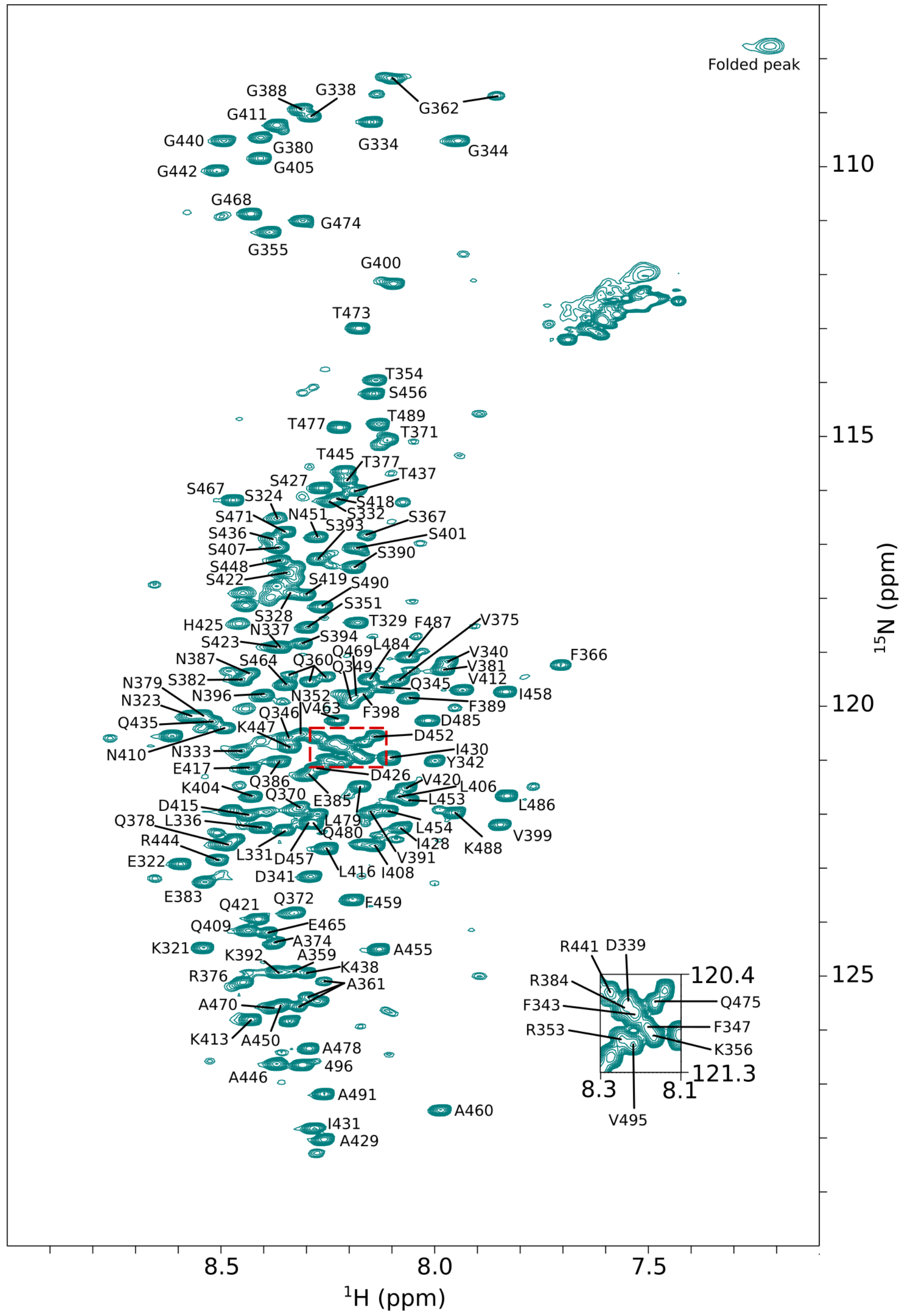


**Supplementary Figure 1. Assignment of Dab2$_{320\text{-}495}$.** The $^{1}$H-$^{15}$N HSQC spectrum of Dab2$_{320\text{-}495}$ showing the backbone resonance assignments as one letter amino acid code. The region marked with red dashed lines is shown as a zoom inside the spectrum. Some residues, e.g. G362, give rise to multiple resonances likely due to proline cis/trans isomerization. Only the main state is uploaded to BMRB (52613).

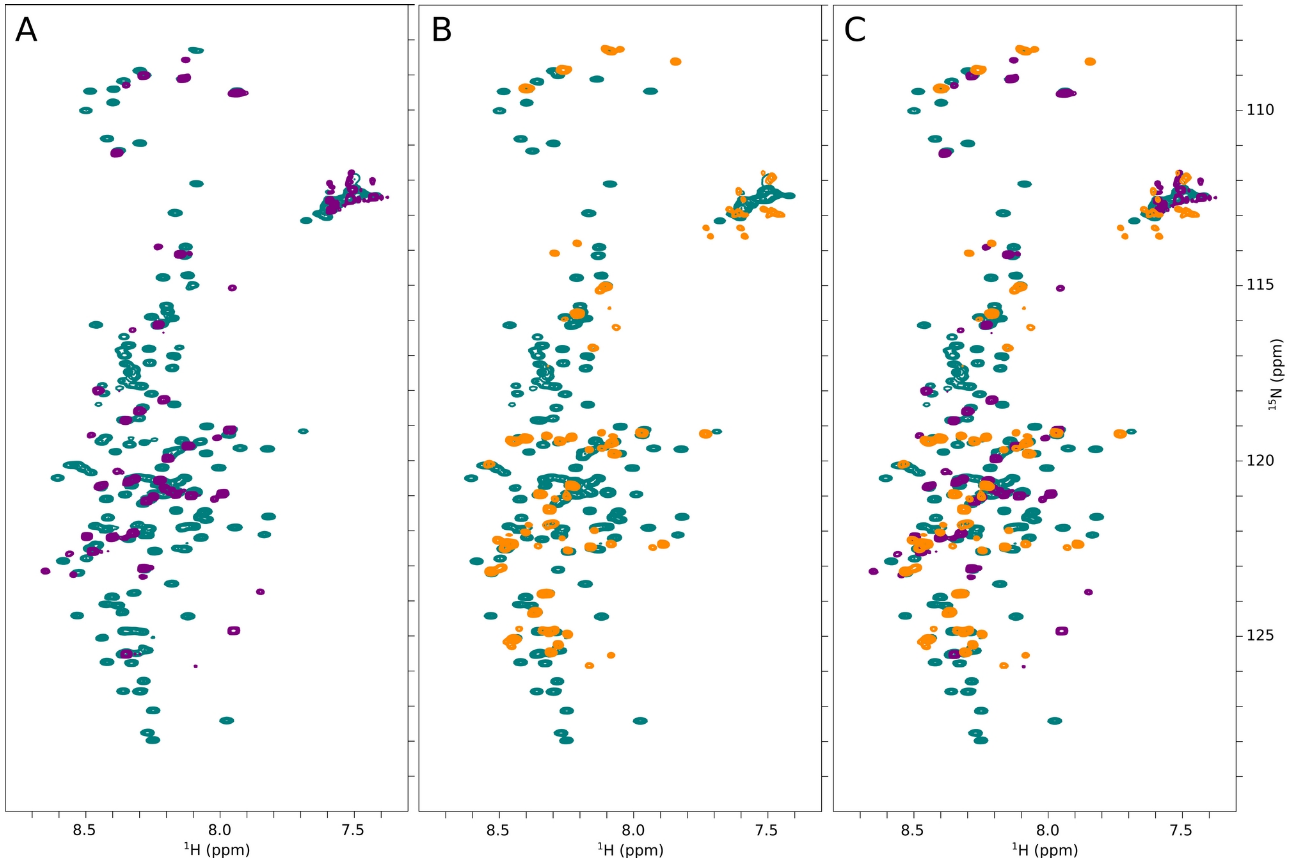


**Supplementary Figure 2. Overlay of $^{1}$H-$^{15}$N HSQC spectra of Dab2$_{320\text{-}495}$ (teal), Dab2$_{328\text{-}360}$ (purple) and Dab2$_{358\text{-}390}$ (orange). (A)** Dab2$_{328\text{-}360}$ spectrum overlaid onto the spectrum of Dab2$_{320\text{-}495}$. **(B)** Dab2$_{358\text{-}390}$ spectrum overlaid onto the spectrum of Dab2$_{320\text{-}495}$. **(C)** Spectra of Dab2$_{328\text{-}360}$ and Dab2$_{358\text{-}390}$ overlaid onto the spectrum of Dab2$_{320\text{-}495}$.

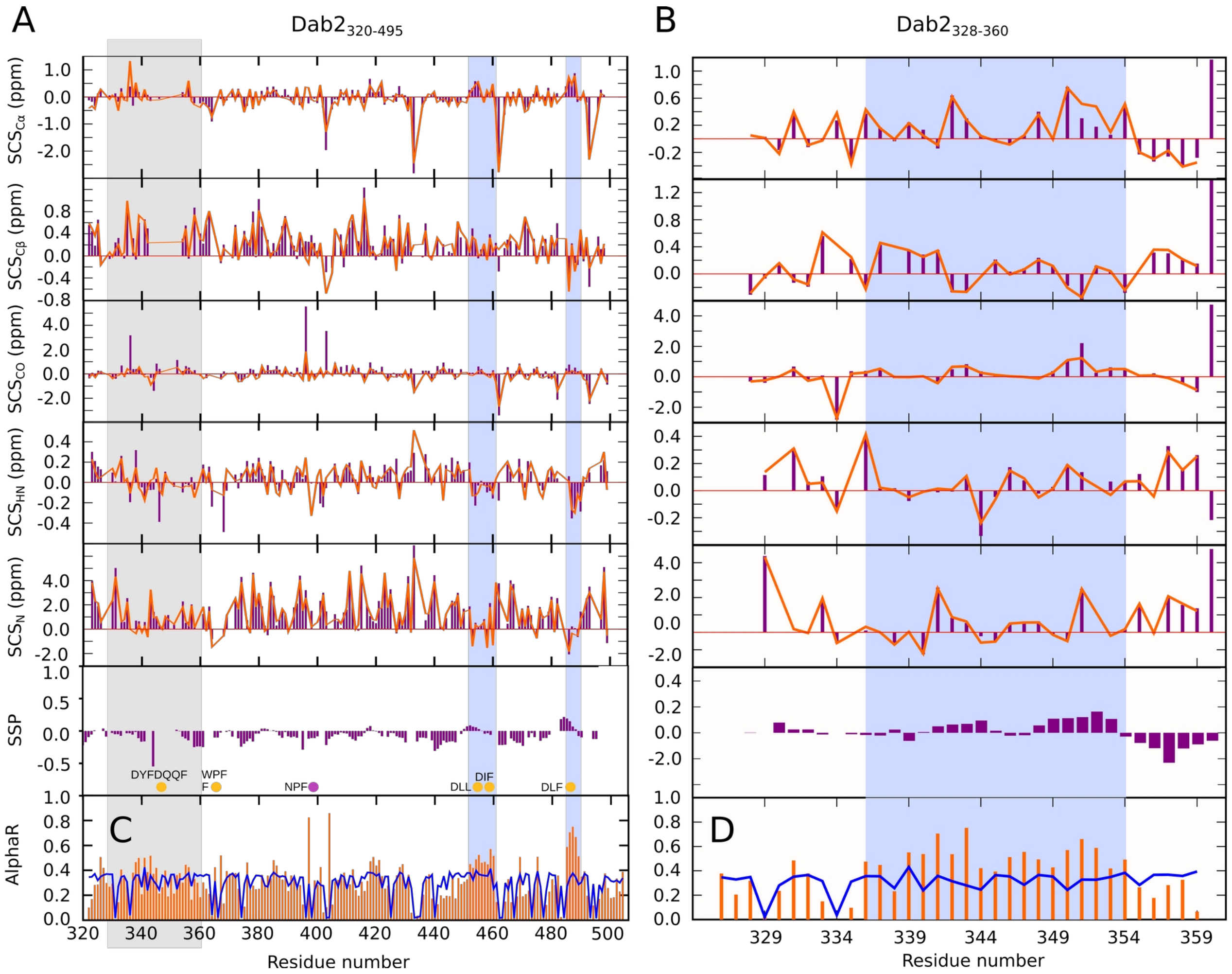


**Supplementary Figure 3. Secondary chemical shifts and secondary structure propensities of (A) Dab2$_{320\text{-}495}$ and (B) Dab2$_{328\text{-}360}$.** SCSs (purple bars) were calculated with respect to random coil chemical shifts from RefDB[54]. Secondary chemical shifts (SSPs)[29] (purple bars) were calculated based on Cα and Cβ chemical shifts. An SSP value of 1 reflects a fully formed helix, a value of -1 reflects a fully extended (β-strand) conformation. Back calculation of SCSs and SSPs on the basis of the ASTEROIDS ensemble is shown as orange lines above the experimental values in purple. An ASTEROIDS ensemble of Dab2$_{320\text{-}495}$ **(C)** and Dab2$_{328\text{-}360}$ **(D)** illustrates a mild increase in helical conformation (AlphaR, $\phi < 0°$, $-120° < \psi < 50°$)[63] (orange bars) as compared to random coil (blue line), from residues 338 to 358 approximately and additional helical sampling around residues 450 to 455 as well as around 480. Regions with increased helical propensity are highlighted in blue. The protein region in Dab2$_{320\text{-}495}$ for which the Dab2$_{328\text{-}360}$ construct was generated is highlighted in gray in panels A and C.

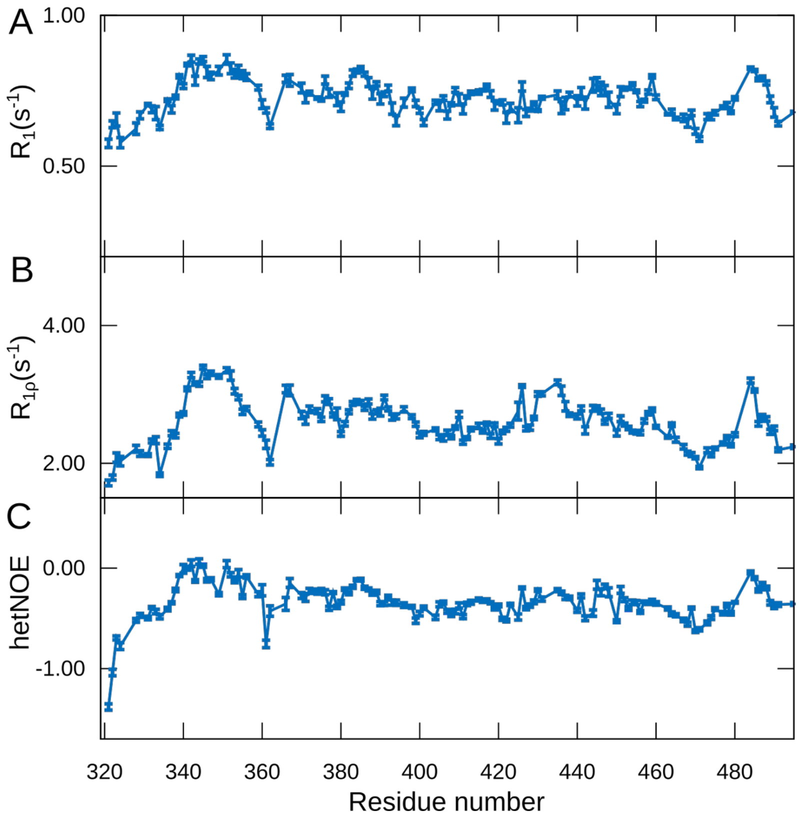


**Supplementary Figure 4. $^{15}$N backbone relaxation of Dab2$_{320\text{-}495}$. (A)** $^{15}$N $R_1$ relaxation, **(B)** $^{15}$N $R_{1\rho}$ spin relaxation, **(C)** {$^{1}$H}-$^{15}$N HetNOE of Dab2$_{320\text{-}495}$. The experiments were recorded on a 270 µM Dab2$_{320\text{-}495}$ sample at a $^{1}$H frequency of 600 MHz.

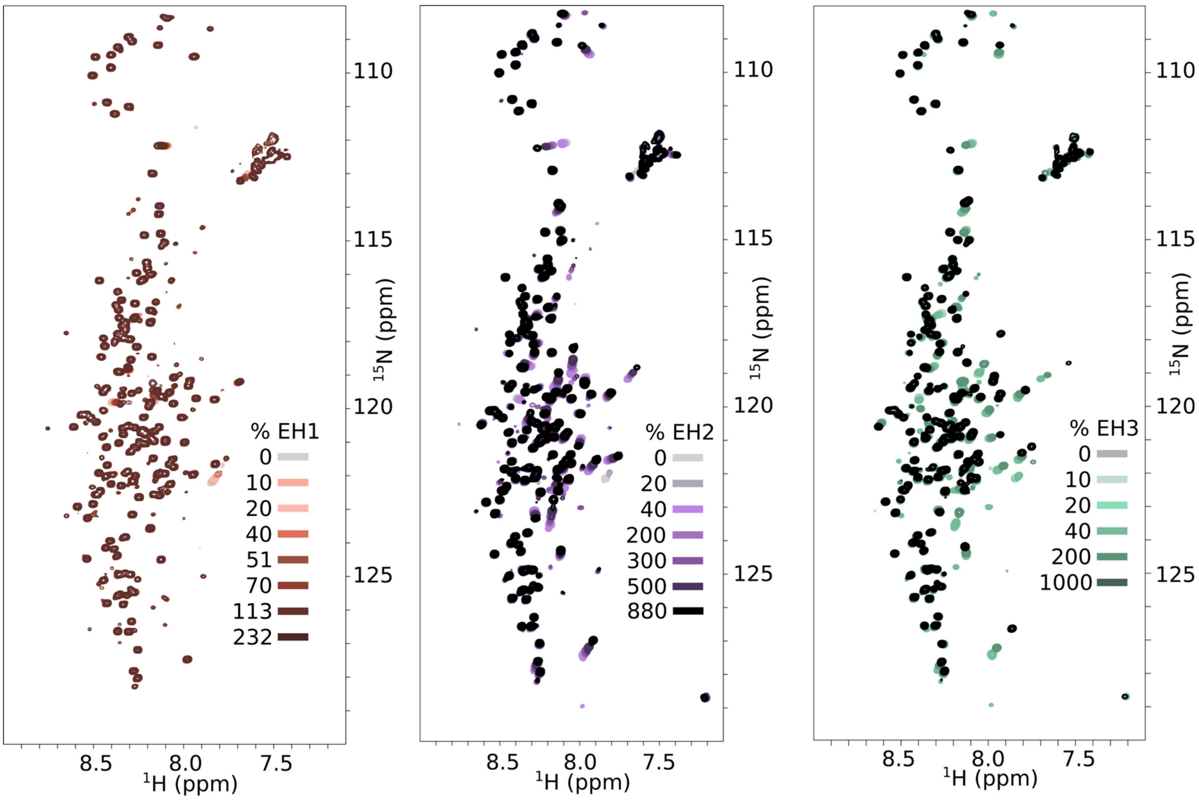


**Supplementary Figure 5. Interaction of $^{15}$N Dab2$_{320\text{-}495}$ with EH1, EH2, and EH3.** $^{1}$H-$^{15}$N HSQC spectra of Dab2$_{320\text{-}495}$ alone and in the presence of increasing concentrations of EH1 (left), EH2 (middle) and EH3 (right). Color codes of the different spectra are indicated in the figure. The concentration of Dab2$_{320\text{-}495}$ was kept constant at 100 µM throughout the titration.

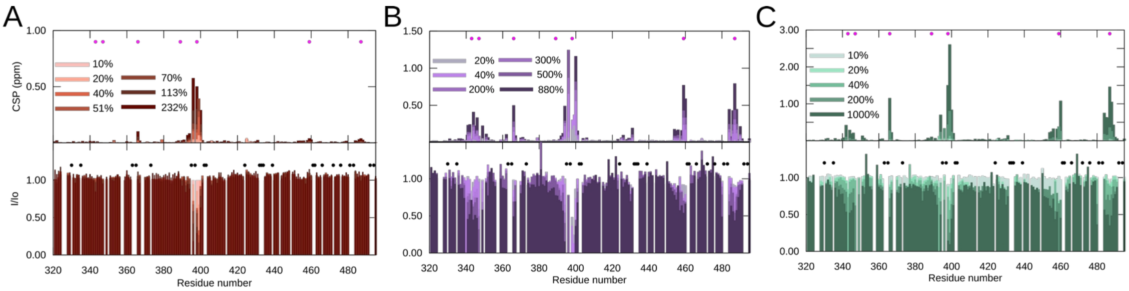


**Supplementary Figure 6. Interaction of Dab2$_{320\text{-}495}$ with the individual EH domains.** Top: CSPs extracted from Dab2$_{320\text{-}495}$ $^{1}$H-$^{15}$N HSQC spectra in the presence of increasing concentrations of EH1 **(A),** EH2 **(B)**, EH3 **(C)** (see also Figure 1D, E, F, respectively). Bottom: Intensity ratio calculated from peak intensities extracted from Dab2$_{320\text{-}495}$ $^{1}$H-$^{15}$N HSQC spectra in the presence of various concentrations of EH1 (A), EH2 (B), EH3 (C) versus the absence of interaction partner. Color legends are displayed in the respective plots. Filled pink circles denote positions of phenylalanines and filled black circles denote positions of prolines.

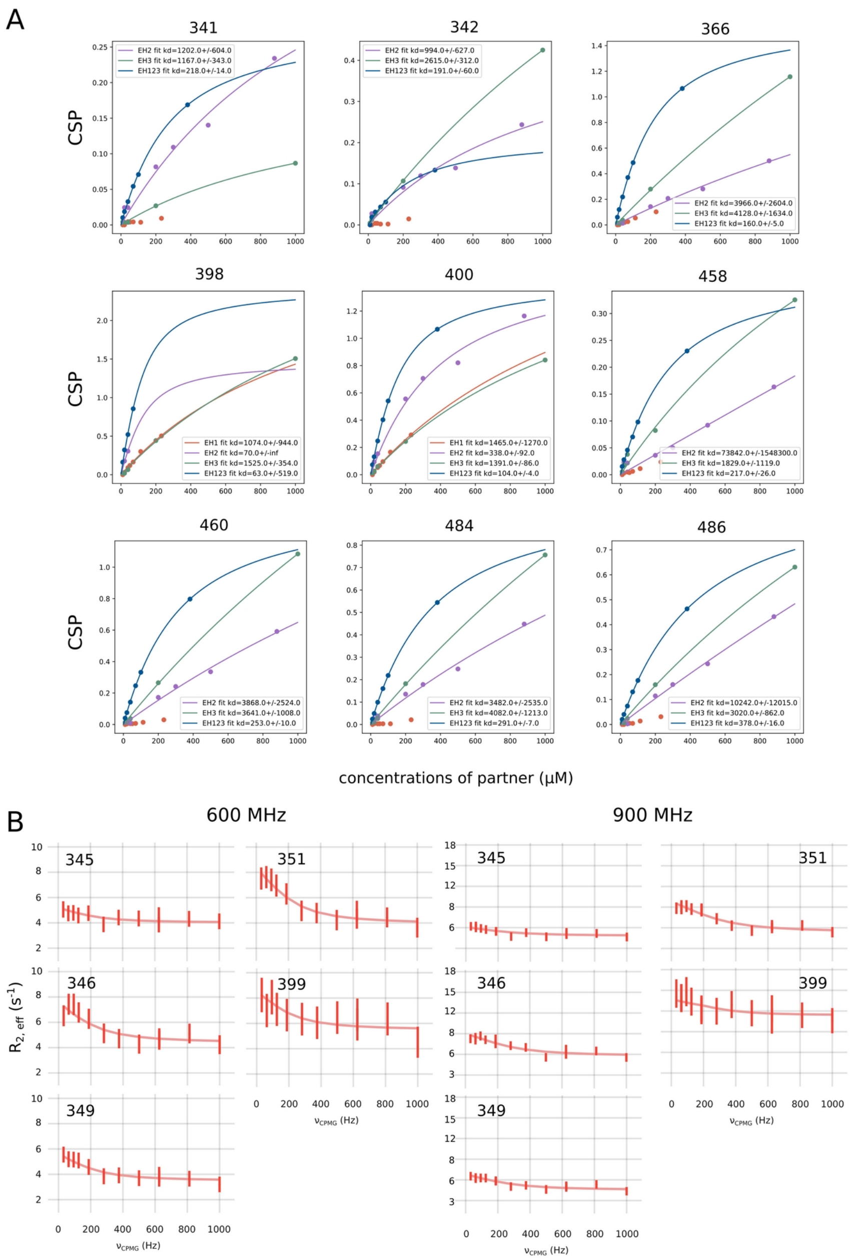


**Supplementary Figure 7. Affinities between the Dab2$_{320\text{-}495}$ NPF motif and EH domains. (A)** CSPs were calculated from $^{1}$H and $^{15}$N chemical shifts at increasing concentrations of EH1, EH2, EH3, or EH123 and plotted against these concentrations (filled points, blue: EH123, brown: EH1, purple: EH2, green: EH3). The data were then fit with a simple binding model (see methods, solid line). Affinities extracted from the fit are displayed next to the color legend in the figure. Units are micromolar. **(B)** CPMG relaxation dispersion curves of five Dab2$_{320\text{-}495}$ residues in the presence of EH2. Dispersion is visible at the NPF motif (residue 399) and around the transient helix at the N-terminus of Dab2$_{320\text{-}495}$ (345-351). The data were fitted with a global fit across 12 residues in the helical region and the NPF motif (residues Val340, Asp341, Gln345, Gln346, Gln349, Ser351, Thr354, Lys356, Phe366, Phe 398, Val399, Asp426) and resulted in an exchange rate $k_{ex}$ of 149 ± 14 $s^{-1}$ and a percentage of bound Dab2$_{320\text{-}495}$ of 3.3 ± 0.3%. The data were acquired at a concentration of 100 μM $Dab2_{320-495}$ with 10% of EH2, resulting in a $K_D$ of 196 μM estimated from the CPMG data and in rough agreement with the chemical shift titrations that suffer from intermediate exchange line broadening around the NPF motif. The data were recorded at $^1H$ frequencies of 600 and 900 MHz.

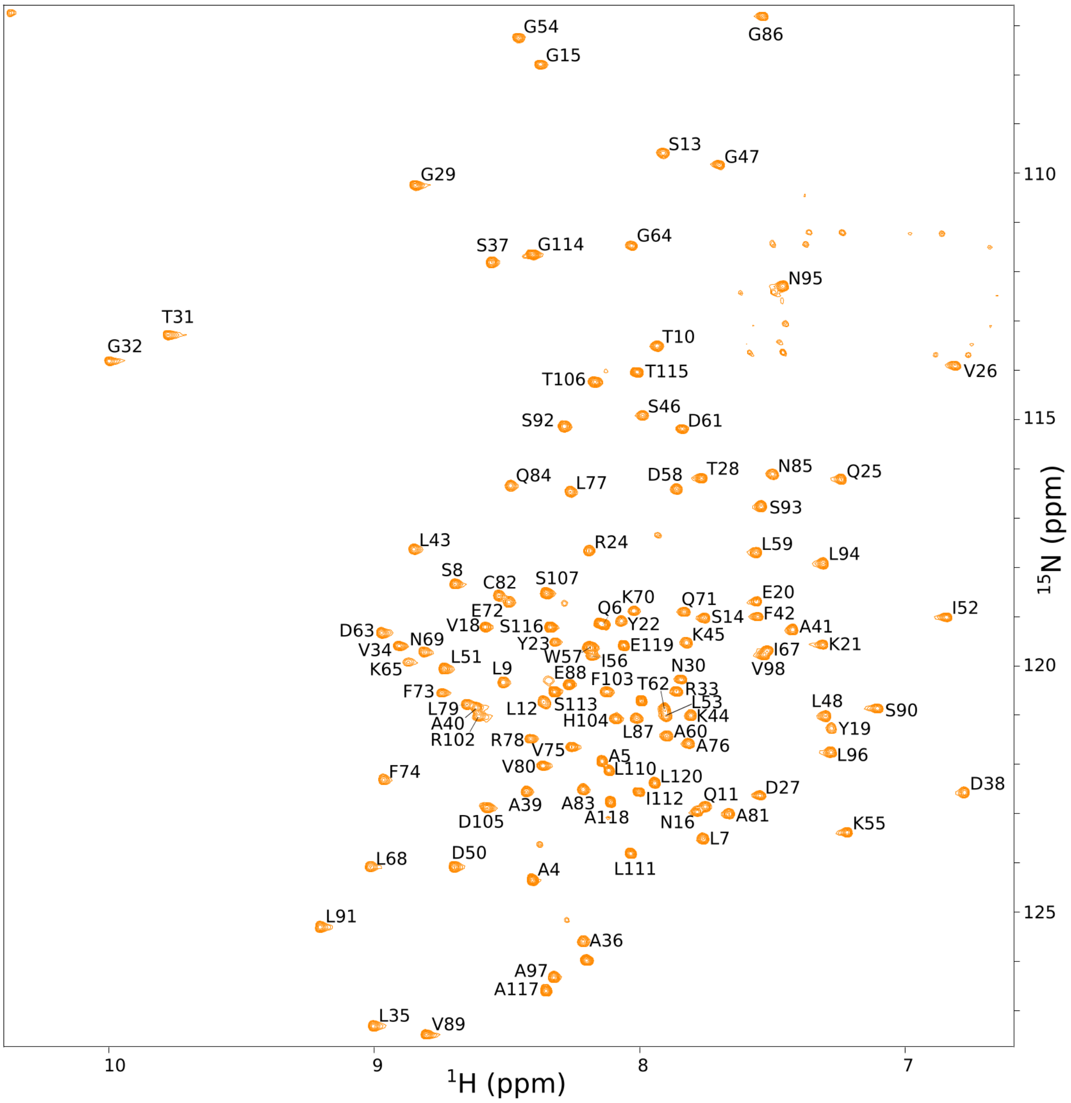


**Supplementary Figure 8. Assignment of EH1.** $^{1}H$-$^{15}N$ TROSY spectrum of EH1, showing the backbone resonance assignments as one letter amino acid code.

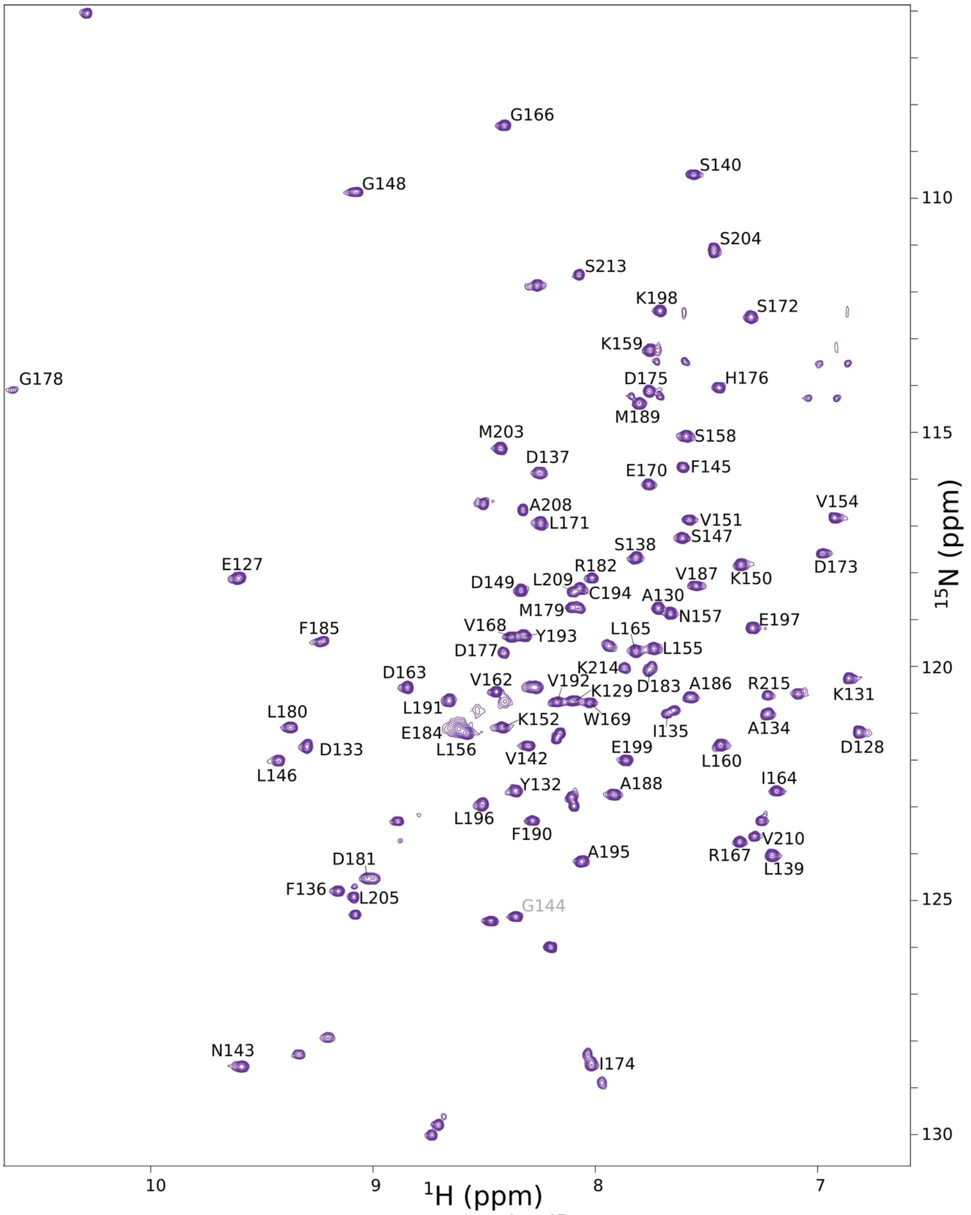


**Supplementary Figure 9. Assignment of EH2.** $^{1}$H-$^{15}$N TROSY spectrum of EH2, showing the backbone resonance assignments as one letter amino acid code. The peak of G144 is folded in the NMR spectrum.

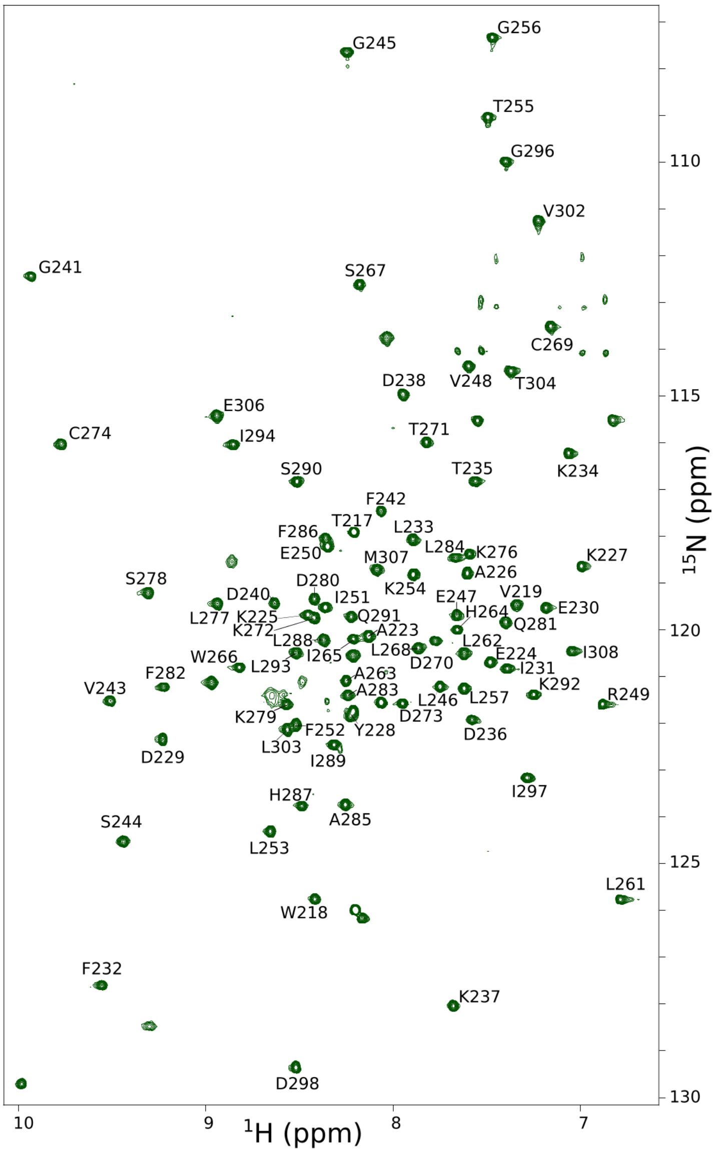


**Supplementary Figure 10. Assignment of EH3.** $^{1}H$-$^{15}N$ TROSY spectrum of EH3, showing the backbone resonance assignments as one letter amino acid code.

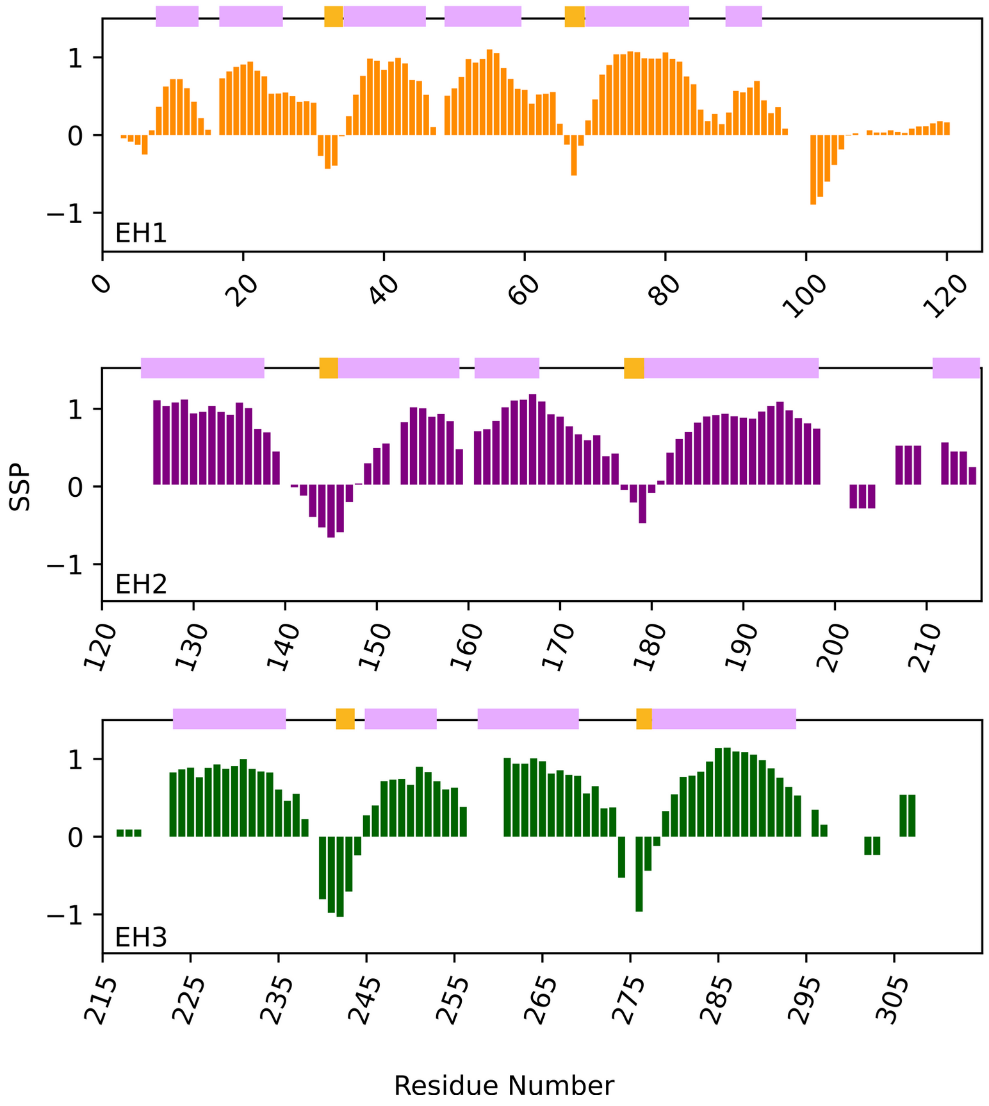


**Supplementary Figure 11. Secondary structure analyses of the individual EH domains.** SSPs of EH1 (top), EH2 (middle) and EH3 (bottom). SSPs were calculated based on Cα and Cβ chemical shifts. A value of 1 reflects a fully formed helix, a value of -1 reflects a fully extended (β-strand) conformation. The pink boxes represent α-helical conformations and the yellow boxes represent β-sheets as observed in the respective PDB structures 1QJT[30] (*mouse* EH1), 1FF1[31] (*human* EH2), 1C07[32] (*human* EH3), showing that the assignments are in agreement with the already published structures.

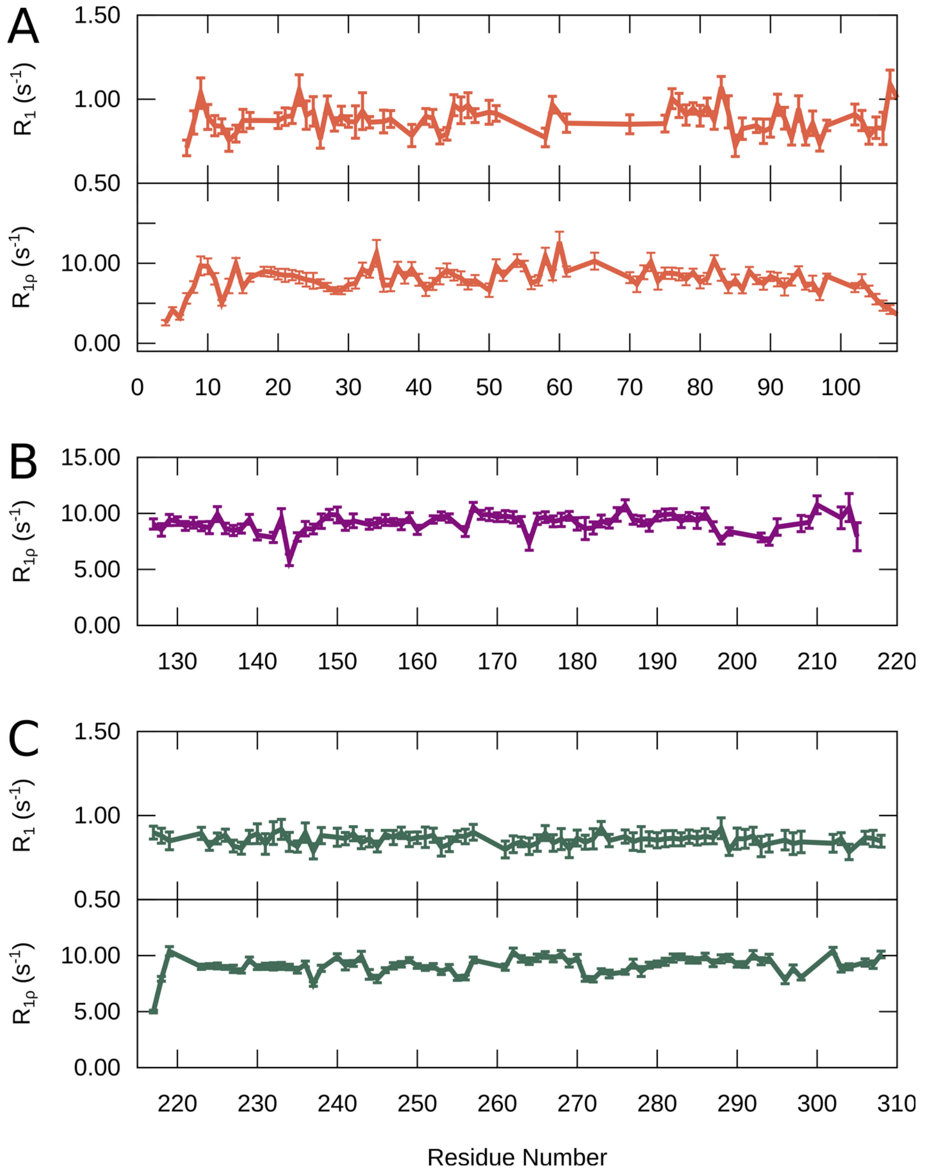


**Supplementary Figure 12. $^{15}$N spin relaxation of EH1, EH2, EH3. (A)** $^{15}$N $R_1$ relaxation and $^{15}$N $R_{1\rho}$ spin relaxation of 100 μM EH1. **(B)** $^{15}$N $R_{1\rho}$ spin relaxation of 100 μM EH2. **(C)** $^{15}$N $R_1$ relaxation of 576 μM of EH3 and $^{15}$N $R_{1\rho}$ spin relaxation of 100 μM EH3. The experiments were all recorded at a $^1$H frequency of 600 MHz.

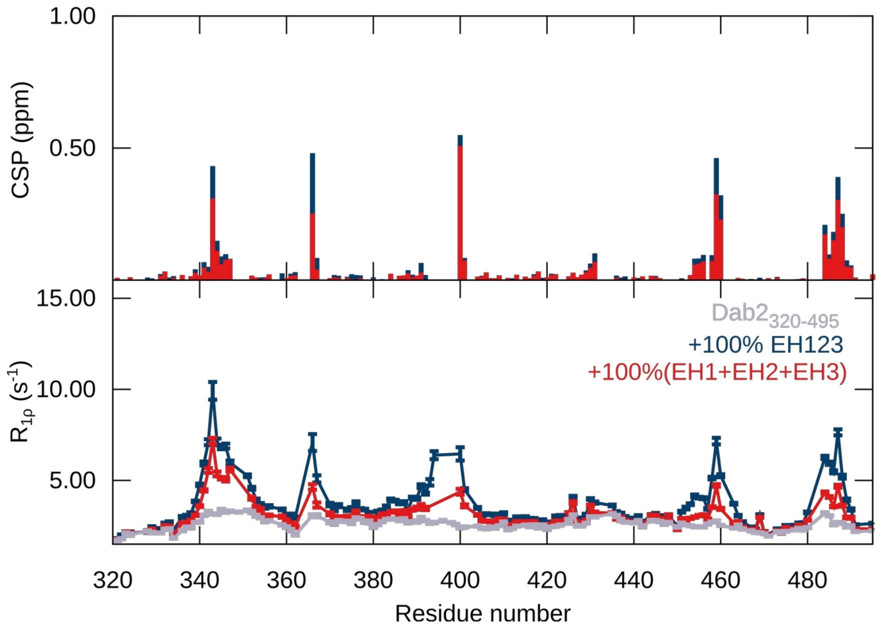


**Supplementary Figure 13. Interaction of Dab2$_{320\text{-}495}$ with EH123 and EH1+EH2+EH3.** CSPs of 100 μM Dab2$_{320\text{-}495}$ in the presence of 100% EH123 (Dark blue) or 100% of each of the individual EH1, EH2, EH3 domains (red). $^{15}$N $R_{1\rho}$ relaxation rates of 100 μM Dab2$_{320\text{-}495}$ alone (gray) and in the presence of 100 μM EH123 (dark blue) or EH1 + EH2 + EH3, each at 100 μM (red).

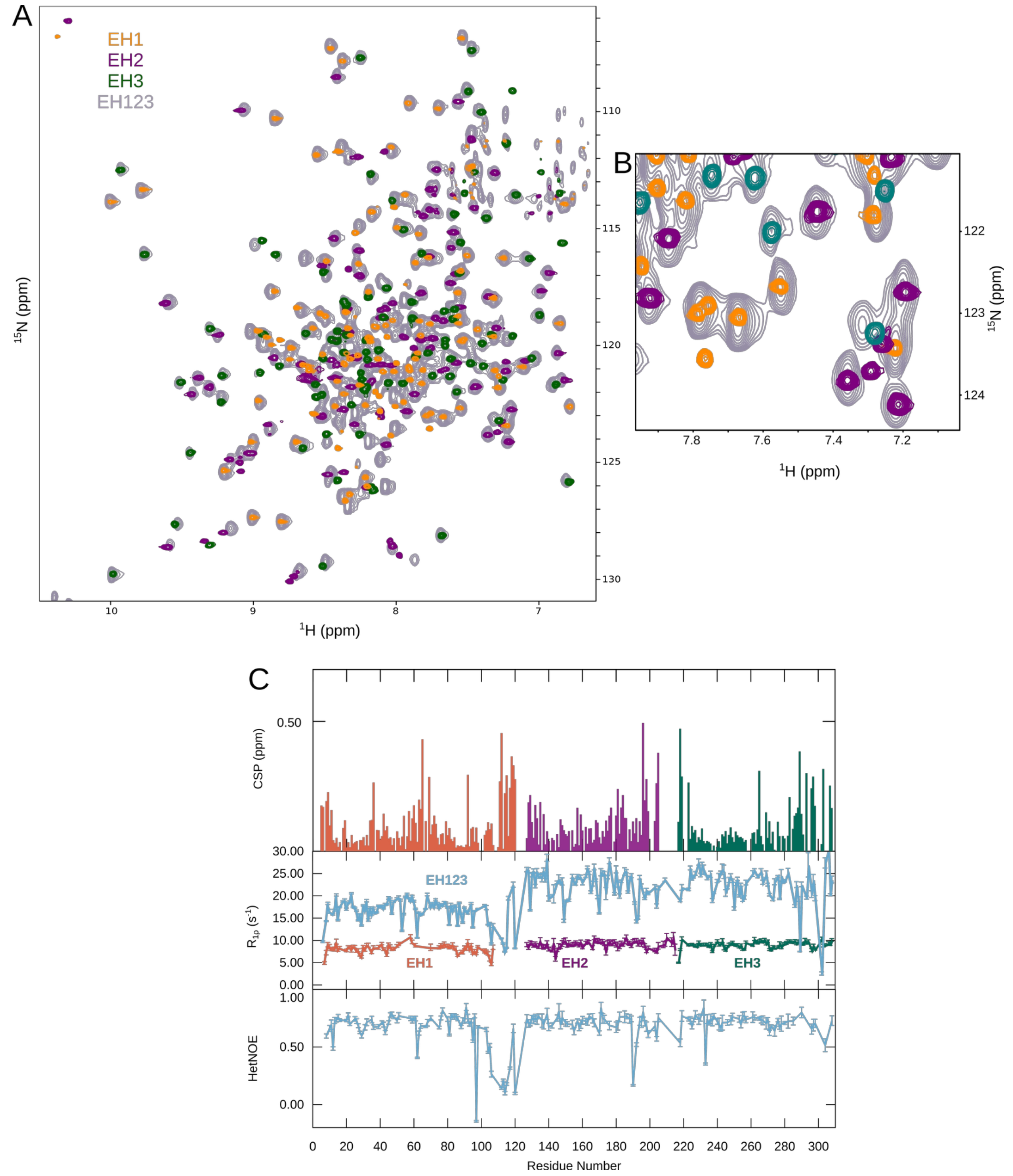


**Supplementary Figure 14. EH2 and EH3 tumble together within EH123. (A)** Overlay of $^{1}$H-$^{15}$N TROSY spectrum of EH123 (gray) and EH1 (orange), EH2 (purple), EH3 (green). **(B)** Zoom into the $^{1}$H-$^{15}$N TROSY spectrum of A. **(C)** CSPs (top) calculated between the $^{1}$H-$^{15}$N TROSY of the $^{15}$N EH123 domain and those of EH1 (orange), EH2 (purple), EH3 (green). $^{15}$N $R_{1\rho}$ spin relaxation (middle) of EH123 (light blue) and EH1 (orange), EH2 (purple), EH3 (green). {$^{1}$H}-$^{15}$N HetNOE (bottom) of EH123 (light blue). The experiments were recorded at a $^{1}$H frequency of 600 MHz. The concentration of EH123 is 260 μM in the standard NMR buffer.

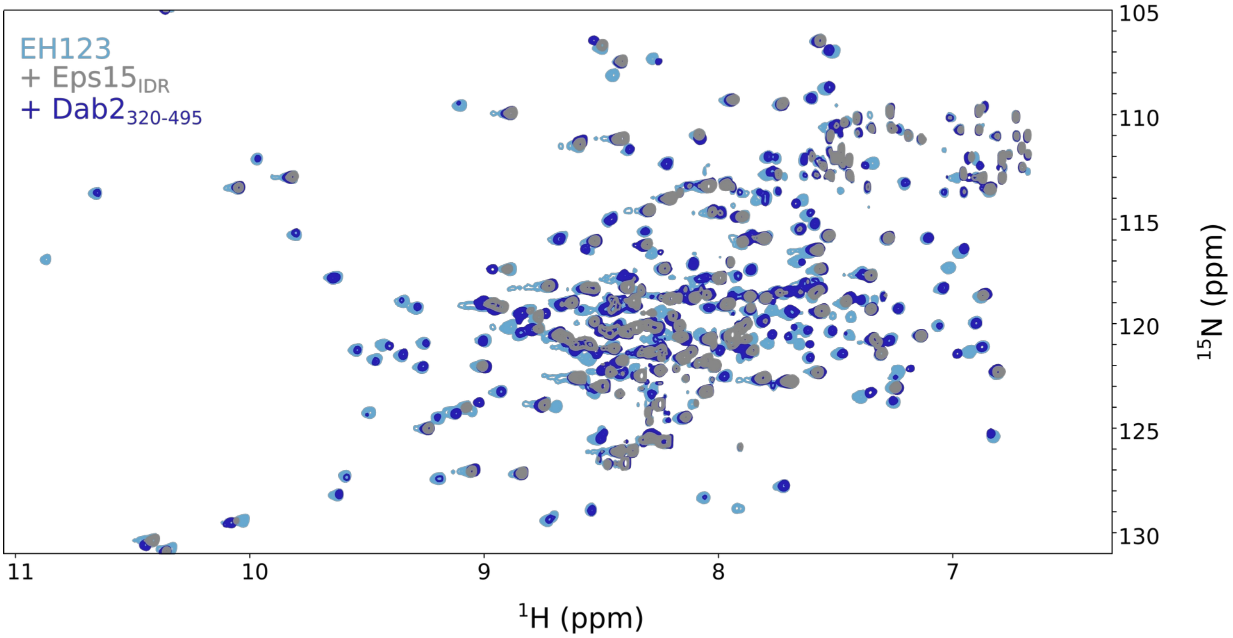


**Supplementary Figure 15. Interaction between $^{15}$N EH123 and Eps15$_{IDR}$ or Dab2$_{320-495}$.** $^{1}$H-$^{15}$N TROSY spectra of 200 µM EH123 (light blue) alone and in the presence of 200 µM Eps15$_{IDR}$ (gray) or 200 µM Dab2$_{320-495}$ (dark blue).

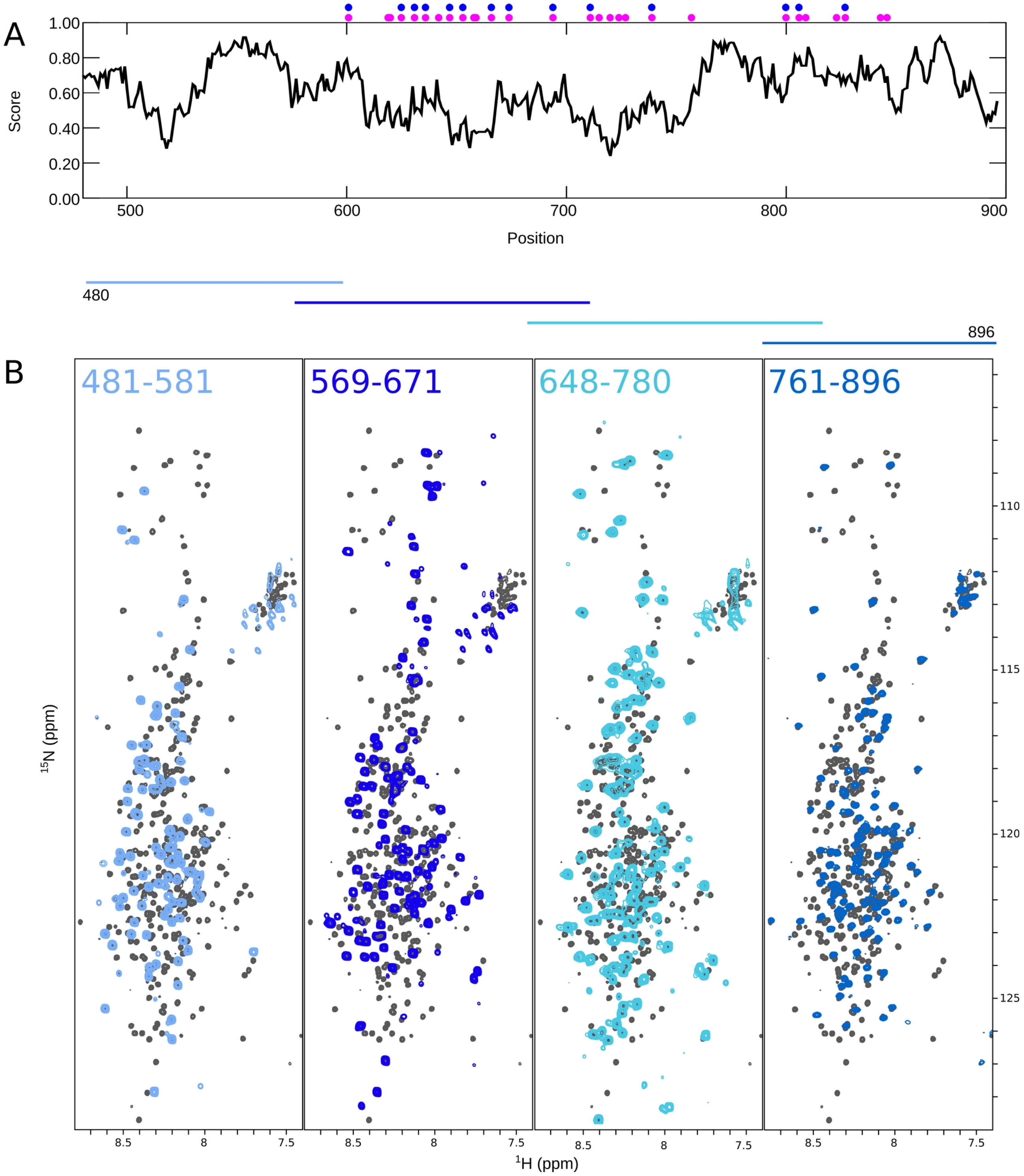


**Supplementary Figure 16. Eps15$_{IDR}$ disorder prediction and NMR spectra. (A)** IUPred2[23] Disorder prediction of Eps15$_{IDR}$ (black) Larger than 0.5: predicted disordered, Smaller than 0.5: predicted ordered. Filled pink circles denote positions of phenylalanines and filled blue circles denote positions of DPF motifs. **(B)** Superimposition of a $^{1}H$-$^{15}N$ HSQC spectrum of Eps15$_{IDR}$ (dark gray) with spectra of the 4 smaller Eps15$_{IDR}$ stretches (different shades of blue).

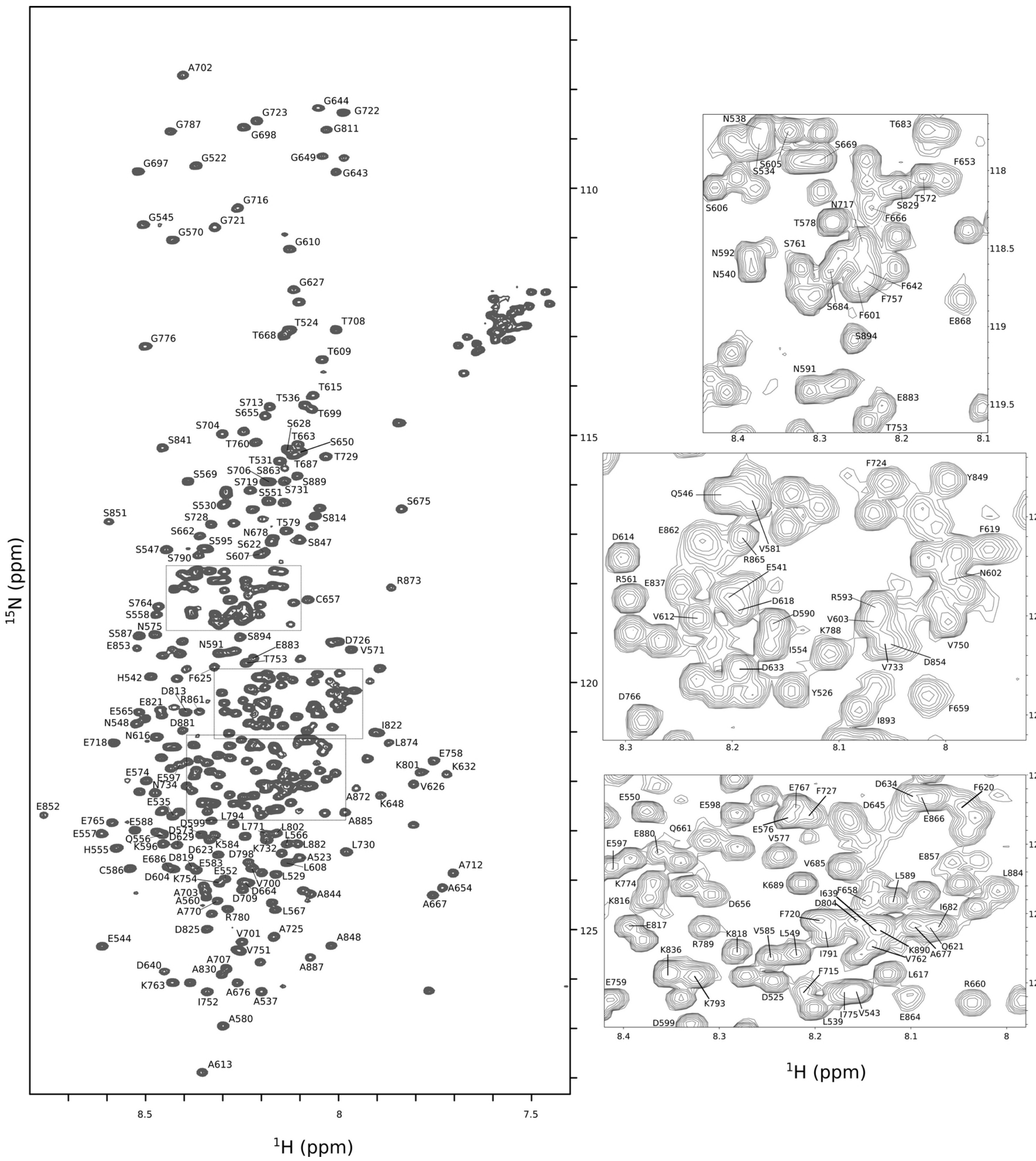


**Supplementary Figure 17. Assignment of Eps15$_{IDR}$.** $^{1}$H-$^{15}$N HSQC spectrum of Eps15$_{IDR}$ showing the backbone resonance assignments as one letter amino acid code. The regions marked with black boxes are shown as zoom-ins next to the spectrum. A702 appears as a spectrally folded peak.

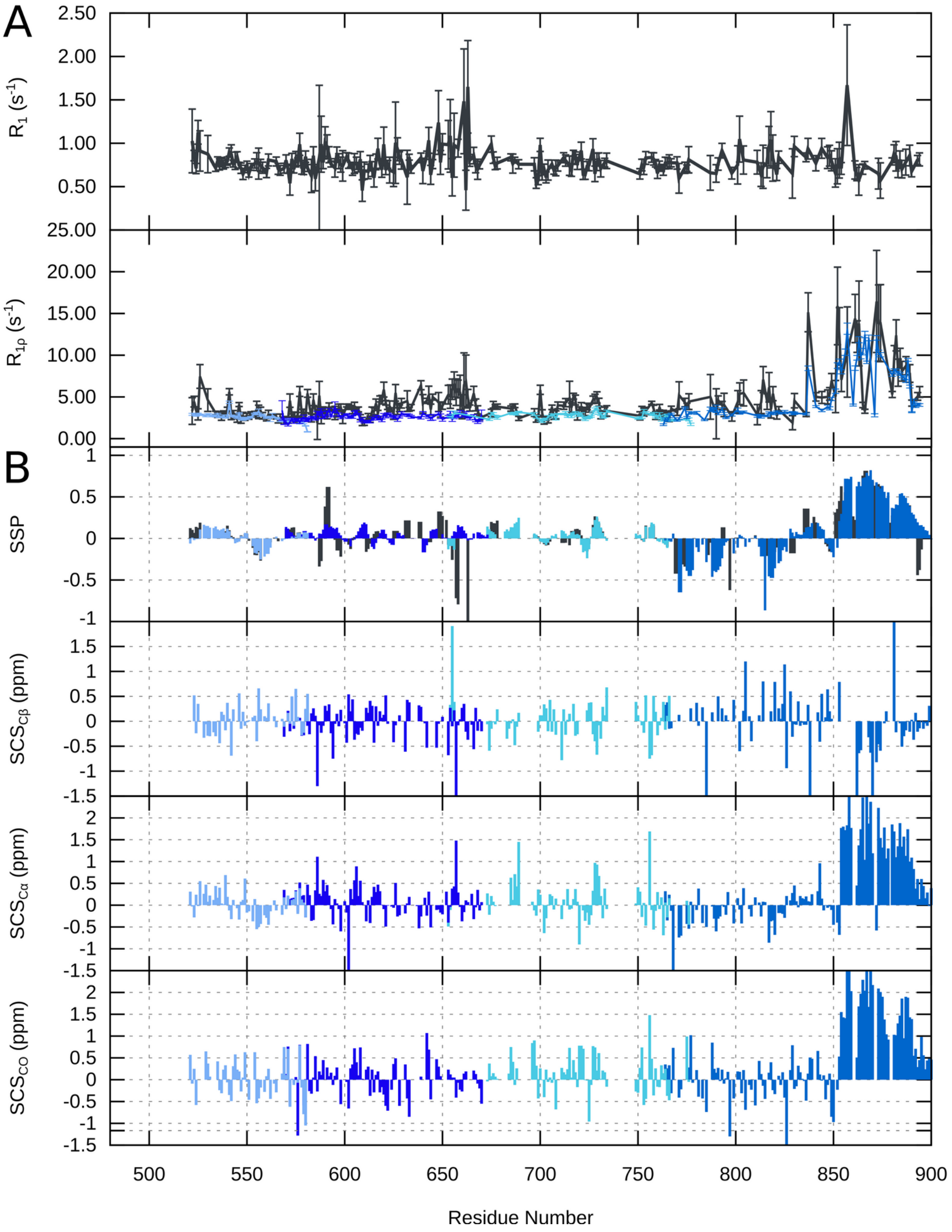


**Supplementary Figure 18. Relaxation and secondary chemical shifts of Eps15$_{IDR}$. (A)** $^{15}N$ $R_1$ and $^{15}N$ $R_{1\rho}$ spin relaxation of Eps15$_{IDR}$ (black) overlaid with the different smaller stretches of Eps15$_{IDR}$ (different shades of blue). **(B)** SSPs[29] of Eps15$_{IDR}$ (black) as well as SSPs and SCSs of Eps15$_{IDR\ 481\text{-}581}$, Eps15$_{IDR\ 569\text{-}671}$, Eps15$_{IDR\ 648\text{-}780}$, and Eps15$_{IDR\ 761\text{-}896}$ (different shades of blue), showing an alpha helical element in the C-terminus (~residues 850-885). SCSs were calculated with respect to random coil chemical shifts, SSPs were calculated based on Cα and Cβ chemical shifts. A value of 1 reflects a fully formed helix, a value of -1 reflects a fully extended (β-strand) conformation.

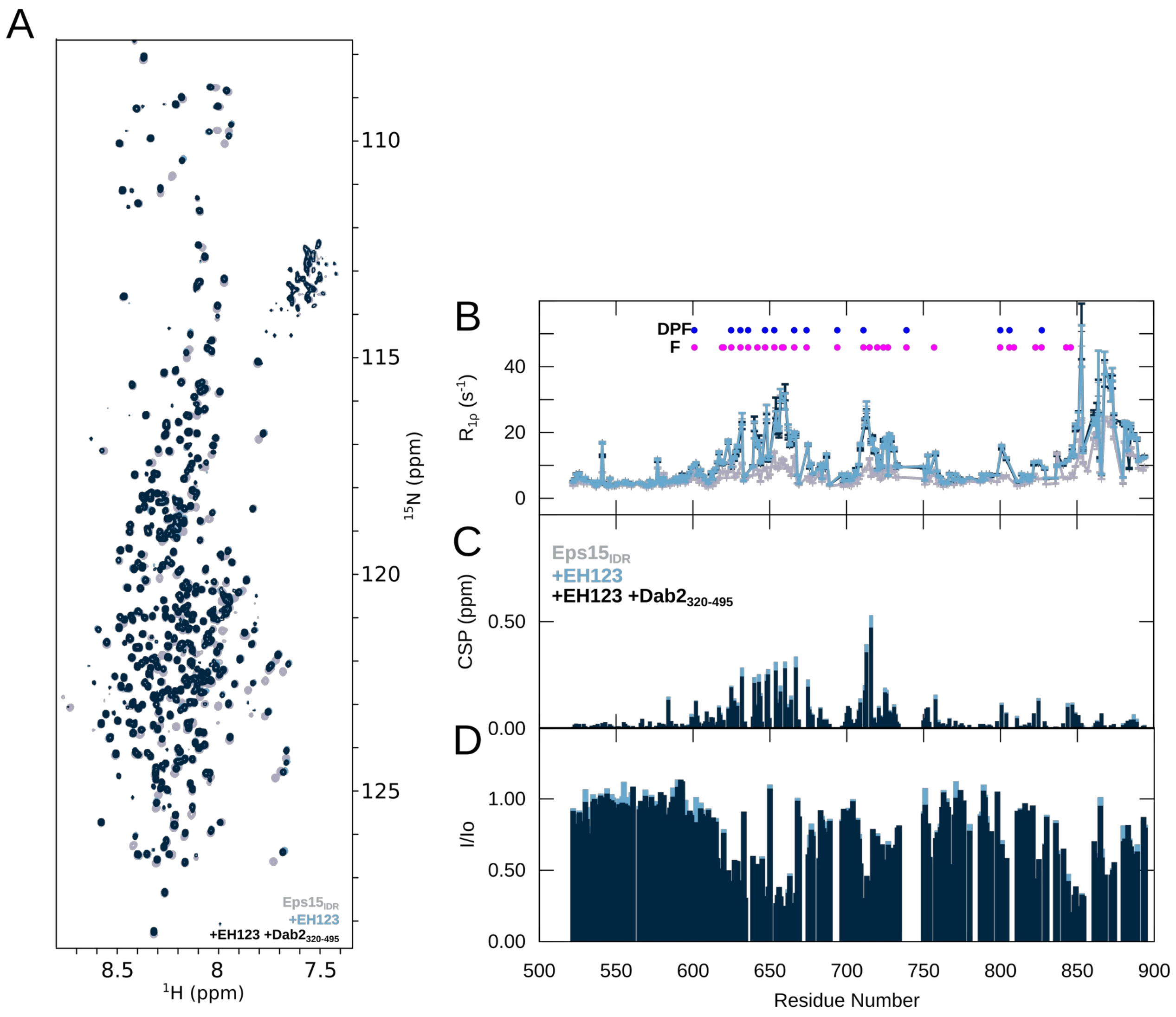


**Supplementary Figure 19. Competition experiment between $^{15}$N Eps15$_{IDR}$ and Dab2$_{320\text{-}495}$ with EH123. (A)** $^{1}$H-$^{15}$N HSQC spectra of 100 µM $^{15}$N Eps15$_{IDR}$ alone and in the presence of 100 µM EH123 as well as both 100 µM EH123 and 100 µM Dab2$_{320\text{-}495}$. **(B)** $^{15}$N $R_{1\rho}$ relaxation rates, **(C)** CSPs, and **(D)** intensity ratios of 100 µM Eps15$_{IDR}$ with 100 µM EH123 or both EH123 and Dab2$_{320\text{-}495}$ at 100 µM each. The color codes are displayed in the respective figure panels.

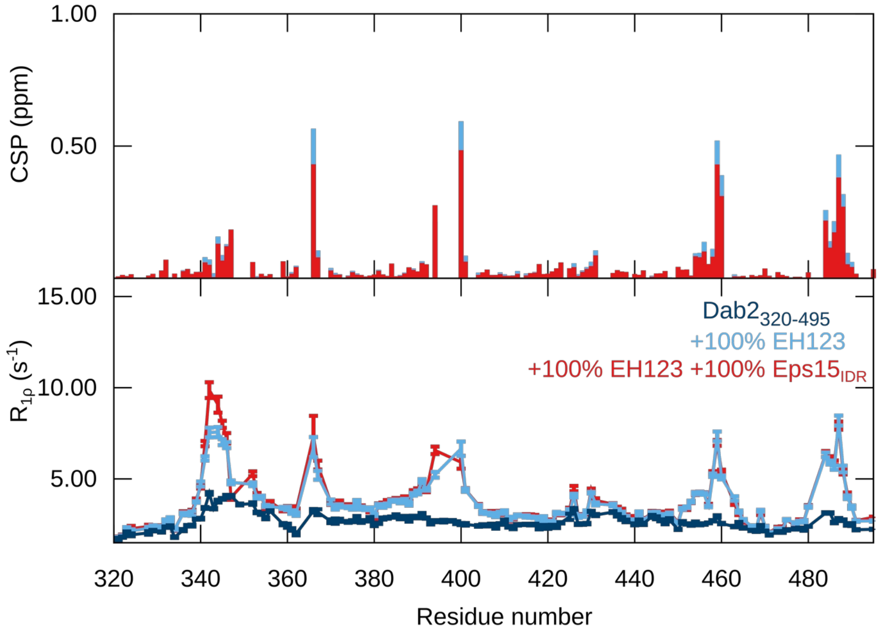


**Supplementary Figure 20. Competition experiment between $^{15}$N Dab2$_{320\text{-}495}$ and Eps15$_{IDR}$ with EH123.** CSPs and $^{15}$N $R_{1\rho}$ relaxation rates of $^{1}$H-$^{15}$N HSQC of $^{15}$N Dab2$_{320\text{-}495}$ (100 µM) with EH123 (100 µM) or both 100 µM EH123 and 100 µM Eps15$_{IDR}$. The color codes are displayed in the plots.

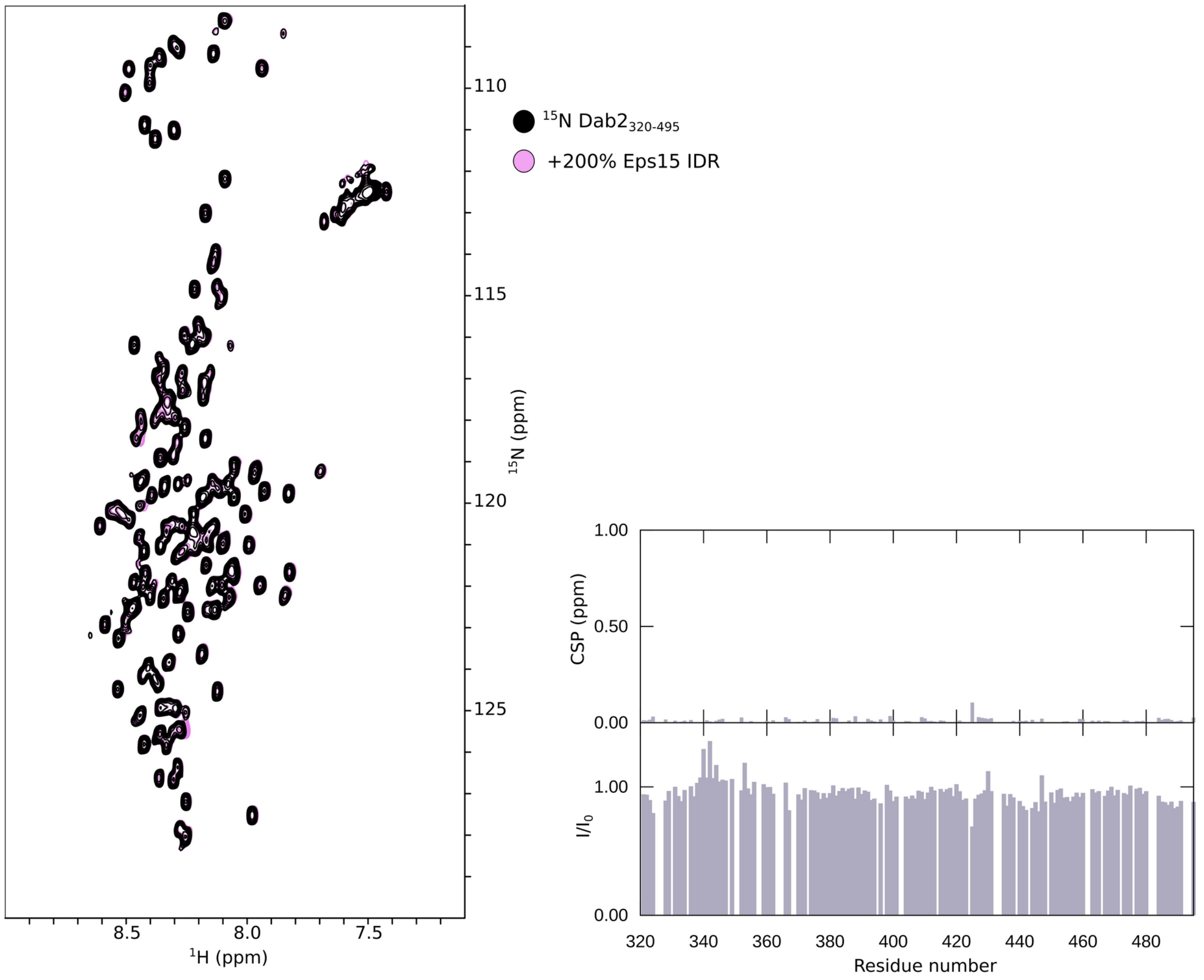


**Supplementary Figure 21. Dab2$_{320\text{-}495}$ and Eps15$_{IDR}$ do not interact.** Overlay of a spectrum of 100 µM Dab2$_{320\text{-}495}$ (black) with that of 100 µM Dab2$_{320\text{-}495}$ + 200% Eps15$_{IDR}$ (pink), as well as the calculated CSPs and intensity ratios based on the two spectra, confirming that the two IDRs do not interact.
